# Disentangling selection on genetically correlated polygenic traits using whole-genome genealogies

**DOI:** 10.1101/2020.05.07.083402

**Authors:** Aaron J. Stern, Leo Speidel, Noah A. Zaitlen, Rasmus Nielsen

## Abstract

We present a full-likelihood method to estimate and quantify polygenic adaptation from contemporary DNA sequence data. The method combines population genetic DNA sequence data and GWAS summary statistics from up to thousands of nucleotide sites in a joint likelihood function to estimate the strength of transient directional selection acting on a polygenic trait. Through population genetic simulations of polygenic trait architectures and GWAS, we show that the method substantially improves power over current methods. We examine the robustness of the method under uncorrected GWAS stratification, uncertainty and ascertainment bias in the GWAS estimates of SNP effects, uncertainty in the identification of causal SNPs, allelic heterogeneity, negative selection, and low GWAS sample size. The method can quantify selection acting on correlated traits, fully controlling for pleiotropy even among traits with strong genetic correlation (|*r_g_*| = 80%; c.f. schizophrenia and bipolar disorder) while retaining high power to attribute selection to the causal trait. We apply the method to study 56 human polygenic traits for signs of recent adaptation. We find signals of directional selection on pigmentation (tanning, sunburn, hair, *P*=5.5e-15, 1.1e-11, 2.2e-6, respectively), life history traits (age at first birth, EduYears, *P*=2.5e-4, 2.6e-4, respectively), glycated hemoglobin (HbA1c, *P*=1.2e-3), bone mineral density (*P*=1.1e-3), and neuroticism (*P*=5.5e-3). We also conduct joint testing of 137 pairs of genetically correlated traits. We find evidence of widespread correlated response acting on these traits (2.6-fold enrichment over the null expectation, *P*=1.5e-7). We find that for several traits previously reported as adaptive, such as educational attainment and hair color, a significant proportion of the signal of selection on these traits can be attributed to correlated response, *vs* direct selection (*P*=2.9e-6, 1.7e-4, respectively). Lastly, our joint test uncovers antagonistic selection that has acted to increase type 2 diabetes (T2D) risk and decrease HbA1c (*P*=1.5e-5).

## Introduction

Genome-wide association studies (GWAS) have shown that many human complex traits, spanning anthropometric, behavioral, metabolic, and many other domains, are highly polygenic.^1–3^ GWAS findings have strongly indicated the action of purifying and/or stabilizing selection acting pervasively on complex traits.^4–7^ Some work has also utilized biobank data to measure the fitness effects of phenotypes using direct measurements of reproductive success.^8^ However, there are few, if any, validated genomic signals of directional polygenic adaptation in humans.

Several factors have contributed to this uncertainty. Chief among them, polygenicity can allow adaptation to occur rapidly with extremely subtle changes in allele frequencies.^9^ Classic population genetics-based methods to detect adaptation using nucleotide data have historically been designed to detect selective sweeps with strong selection coefficients, unlikely to occur under polygenic architecture.^10^ Polygenic adaptation, after a shift in the fitness optimum, can occur rapidly while causal variants only undergo subtle changes in allele frequency.^11^ After a transient period during which the mean of the trait changes directionally, a new optimum is reached and the effect of selection will then largely be to reduce the variance around the mean.^12^ However, identifying the genomic footprints of the transient period of directional selection is of substantial interest because it provides evidence of adaptation.

To this end, the advent of GWAS has ushered in a series of methods which take advantage of the availability of allele effect estimates by aggregating subtle signals of selection across association-tested loci. For example, some methods (e.g., the *Q_x_*test) compare differences in population-specific polygenic scores -- an aggregate of allele frequencies and allele effect estimates -- across populations, and tests whether they deviate from a null model of genetic drift.^13^ Other methods have generalized this test, e.g. to identify and map polygenic adaptations to branches of an admixture graph.^14^ Whereas the aforementioned methods exploit between-population differentiation to detect polygenic adaptation, another class of methods is based on within-population variation. For example, selection scans based on singleton density score (SDS) have demonstrated utility in detecting polygenic adaptation via the correlation of SNPs’ effect estimates and their SDSs.^15^ Another test looks for dependence of derived allele frequencies (DAF) on SNP effect estimates.^16^

Several powerful tests for selection were developed to take advantage of recent advances in ancestral recombination graph (ARG^17^) and whole-genome genealogy inference. Such methods enjoy better power in detecting selection as the ARG, if observed directly, fully summarizes the effects of selection on linked nucleotide data. We note that several methods, notably *ARGweaver*^18^ infer the strictly-defined ARG; by contrast, methods such as *Relate*^19^ infer a series of trees summarizing ancestral histories spanning chunks of the genome. For example, the % test estimates changes in the population mean polygenic score over time by using the coalescent tree at a polymorphic site as a proxy for its allele frequency trajectory; the sum of these trajectories weighted by corresponding allelic effect estimates forms an estimate of the polygenic score’s trajectory^20^. Speidel, *et al*. (2019) also designed non-parametric test for selection based on coalescence rates of derived- and ancestral-allele-carrying lineages calculated empirically from the coalescent tree inferred by *Relate*.^19^ However, these methods ultimately treat the coalescent tree as a fixed, observed variable, where it is actually hidden and highly uncertain. Furthermore, most methods infer the tree under a neutral model, and thus provide biased estimates of the genealogy under selection.

To address these issues, we recently developed a full-likelihood method, CLUES, to test for selection and estimate allele frequency trajectories.^21^ The method works by stochastically integrating over both the latent ARG using Markov Chain Monte Carlo, and the latent allele frequency trajectory using a dynamic programming algorithm, and then using importance sampling to estimate the likelihood function of a focal SNP’s selection coefficient, correcting for biases in the ARG due to sampling under a neutral model.

Beyond the issue of statistical power, tests for polygenic adaptation can in general be biased when they rely on GWAS containing uncorrected stratification. For example, several groups found strong signals of height adaptation in Europe^13–15,22–24^; later, it was shown that summary statistics from the underlying meta-analysis (GIANT, a.k.a. Genetic Investigation of ANthropometric Traits) were systematically biased due to uncorrected stratification, and subsequent tests for selection on height failed to be reproduced using properly corrected summary statistics^20,25,26^. However, beyond this case, the extent to which other signals of polygenic selection may be inflated by uncorrected stratification is an open question. Here, we investigate the robustness of the new likelihood method to uncorrected stratification and compare it to another state-of-the-art method (tSDS), showing that our likelihood method is less prone to bias but has substantially improved power.

Another issue faced by current methods to detect polygenic adaptation is confounding due to pleiotropy. For example, direct selection on one trait may cause a false signal of selection on another, genetically-correlated trait. While a variant of the *Q_x_* test has been proposed for the purpose of controls for pleiotropy, this method relies of signals of between-population differentiation to test for selection, and is not directly applicable to test multiple traits jointly.^24^

Here, we present a full-likelihood method (Polygenic Adaptation Likelihood Method, PALM) that uses population DNA sequence data and GWAS summary statistics to estimate direct selection acting on a polygenic trait. We demonstrate robustness by exploring potential sources of bias, including uncorrected GWAS stratification. We also introduce a variant on our method which controls for pleiotropy by testing ≥2 traits for selection jointly. We show our method not only fully controls for this bias, but retains high power to distinguish direct selection from correlated response even in traits with strong genetic correlation (up to 80%), and has unique power to detect cases of antagonistic selection on genetically correlated traits. We explore the behavior of the test when traits with causal fitness effects are excluded to illustrate limitations and proper interpretation of these selection and correlated response estimates.

## Model

### Linking SNP effects to selection coefficients

Let *β* be the effect of a SNP on a trait. We model the selection coefficient acting on this SNP using the Lande approximation^27^ *s* ≈ *βω*, where ω is the selection gradient, the derivative of fitness with respect to trait value. If *β* is measured in phenotypic standard deviations, then *ω* is the so-called selection intensity. Chevin and Hospital (2008) showed that for a neutral ‘tag’ SNP with frequency *u* = 1 − *v* and genotypic correlation *r* to a SNP with selection coefficient s, and allele frequencies *p* and *q=1-p*, to a first approximation the linked neutral SNP effectively undergoes selection with 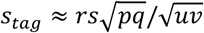.^28^ Applying this principle to the multivariate Lande approximation, we find that *s_tag_ ≈ β_tag_ω*, where 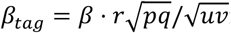 is the marginal effect of the tag SNP, assuming no linkage disequilibrium between the tag SNP and any other causal SNP other.

### Inferring the selection gradient using a full-likelihood model

Our likelihood model builds heavily on Stern, *et al*. (2019), which developed importance sampling approaches for estimating the likelihood function of the selection coefficient acting on a SNP, *L^SNP^* (*s*).^21^ Let *β*_(*i*)_ be the effect of SNP *i* on the trait. Based on these SNP-level selection likelihoods, we model the likelihood function for the selection differential acting on a trait as the product of the SNP likelihoods evaluated at selection coefficients under the Lande approximation:

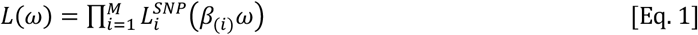

We provide details for calculating this likelihood function using an importance sampling approach based on Stern, *et al*. (2019) (see Methods).^21^ Given this likelihood function, we estimate ωusing its maximum-likelihood estimate. This model is used by our so-called marginal test PALM.

### Fitness effects of multiple traits

To model fitness effects of multiple traits jointly, here we propose a multivariate extension of the Lande approximation which links pleiotropic SNP effects to the selection coefficient. Let *β* be a vector of a particular SNP’s effects on *d* distinct traits. We assume the selection coefficient acting on this SNP follows a multivariate version of the Lande approximation,^27^

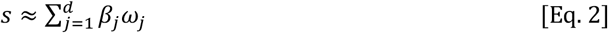

where *ω* now is a vector of selection gradients for each of the *d* traits. The results of Chevin and Hospital (2008) apply directly given this approximation for the selection coefficient, and we now express the likelihood of the selection gradient using Eq. 2: 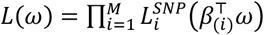. We can solve for the maximum-likelihood estimate of *ω*jointly using standard optimization. This model is used by our joint test J-PALM.

## Results

### Simulations

#### Overview of simulations

We conducted evolutionary simulations of polygenic adaptation acting on a wide range of multi-trait polygenic architectures. Our simulated traits are based on SNP heritability and genetic correlation estimates for 23 real human traits^29,30^; unless otherwise stated, we simulate positive selection on/test for selection on a trait modeled after the heritability of schizophrenia(*h*^2^ = 0.45), and in most of our pleiotropy analyses we used parameters based on schizophrenia and its genetic correlation with 3 other traits: bipolar disorder, major depression, and anorexia. In most of our analysis we refer to these traits as Trait I/II/III/IV (corresponding to models of schizophrenia/bipolar/depression/anorexia, respectively). As our method is based on aggregating population genetic signals of selection with GWAS summary statistics, we also simulated GWAS in samples of modern-day individuals (*N* = 10^5^). Our simulated summary statistics incorporate all of the following sources of bias found in GWAS, unless stated otherwise: random noise in the effect estimates; Winner’s Curse bias in the effect estimates (unless stated otherwise, we ascertain SNPs with associations *P* < 5 × 10^−8^ for at least 1 trait analyzed); uncertainty in the location of the causal SNP (we ascertain the top GWAS hit throughout the linked region); and environmentally correlated noise across traits (only relevant to simulations of pleiotropic architectures). Average selection coefficients, allele frequency changes, and population phenotype changes are detailed in Supp. Tab. 1. Furthermore, we also simulate a number of scenarios which violate our model assumptions, to assess our method’s robustness: these include uncorrected GWAS stratification; purifying/stabilizing selection; underpowered/uneven GWAS sample sizes; and allelic heterogeneity (i.e., multiple linked causal SNPs).

For each causal locus, we simulate haplotype data for a sample of *n* = 400 1Mbp-long chromosomes (mutation and recombination rates *μ = r* = 10^−8^ and effective population size *N_e_* = 10^4^unless stated otherwise), on which we applied *Relate*, a state-of-the-art method for tree inference^19^, to infer the coalescent tree at SNPs ascertained in this GWAS. However, we point out that our approach can be applied to any pre-existing method for estimating/sampling these trees (e.g. *ARGweaver*^18^). We then conduct importance sampling to estimate the likelihood function of the selection gradient – i.e., the effect of a unit increase in phenotypic values on fitness – for individual traits (i.e., estimated marginally), as well as sets of genetically correlated traits (i.e., estimated jointly). Our method, *Polygenic Adaptation Likelihood Method* (PALM), can be used to estimate *ω* for polygenic traits.

#### Improved power to detect selection and estimates of the selection gradient

We ran PALM to test for selection on our simulations of polygenic trait architectures, described above (and in more detail in Appendix). We estimate the selection gradient and standardize this quantity by its standard error, estimated through block-bootstrap, to conduct a Wald test on whether the selection gradient is non-zero.

First, we conducted simulations at different values of the selection gradient, ranging from neutral (*ω* = 0) to strong (*ω* = 0.1, average change of mean phenotype of ~2 standard deviations), and compared the statistical power of PALM to that of tSDS (Fig 1A). Summaries of SNP selection coefficients, allele frequency changes, and phenotypic changes are detailed in Supp. Tab. 1. We simulate 5Mb haplotypes for a trait with polygenicity (i.e., number of causal SNPs) *M* = 1,000; we sample *n* = 178 haplotypes for PALM and *n* = 6,390for tSDS, corresponding to the sample sizes we used in our application to 1000 Genomes British (GBR) individuals *vs* the sample used by Field, *et al*. (2016) from the UK10K. Here we ascertain only causal SNPs, but SNP effects are still estimated through an association test (unless otherwise stated, all other simulations incorporate uncertainty in the causal SNP). Both methods are well calibrated under the null (*ω* = 0, Fig 1A). But we find that despite having a much smaller sample size, PALM has substantially improved power to detect selection at all levels (Fig 1A), especially at weaker values of the selection gradient, where tSDS has essentially no power (*ω* ≤ 0.05). PALM is also capable of estimating the selection gradient (Fig. 1A, Table 1). These estimates are well-calibrated, with empirical standard errors closely matching estimated standard errors, except when the selection gradient is exceptionally strong (*ω* ≥ 0.1) (Table 1).

**Table 1:** Selection gradient estimates and standard errors. Summary statistics for the accuracy and calibration of estimates also used in Figure 1 (see caption for simulation details). Mean s.e. is the mean nominal standard error. Simulations are the same as used in Figure 1A.

| $\omega$ | mean $\hat{\omega}$ | sd($\hat{\omega}$) | MSE($\hat{\omega}$) | Mean se($\hat{\omega}$) |
| --- | --- | --- | --- | --- |
| 0 | 0.0053 | 0.0226 | 0.0232 | 0.0246 |
| 0.025 | 0.0306 | 0.0225 | 0.0232 | 0.0243 |
| 0.05 | 0.0465 | 0.0243 | 0.0245 | 0.0266 |
| 0.075 | 0.0931 | 0.0211 | 0.0278 | 0.023 |
| 0.1 | 0.1223 | 0.0236 | 0.0325 | 0.0255 |

**Figure 1:**
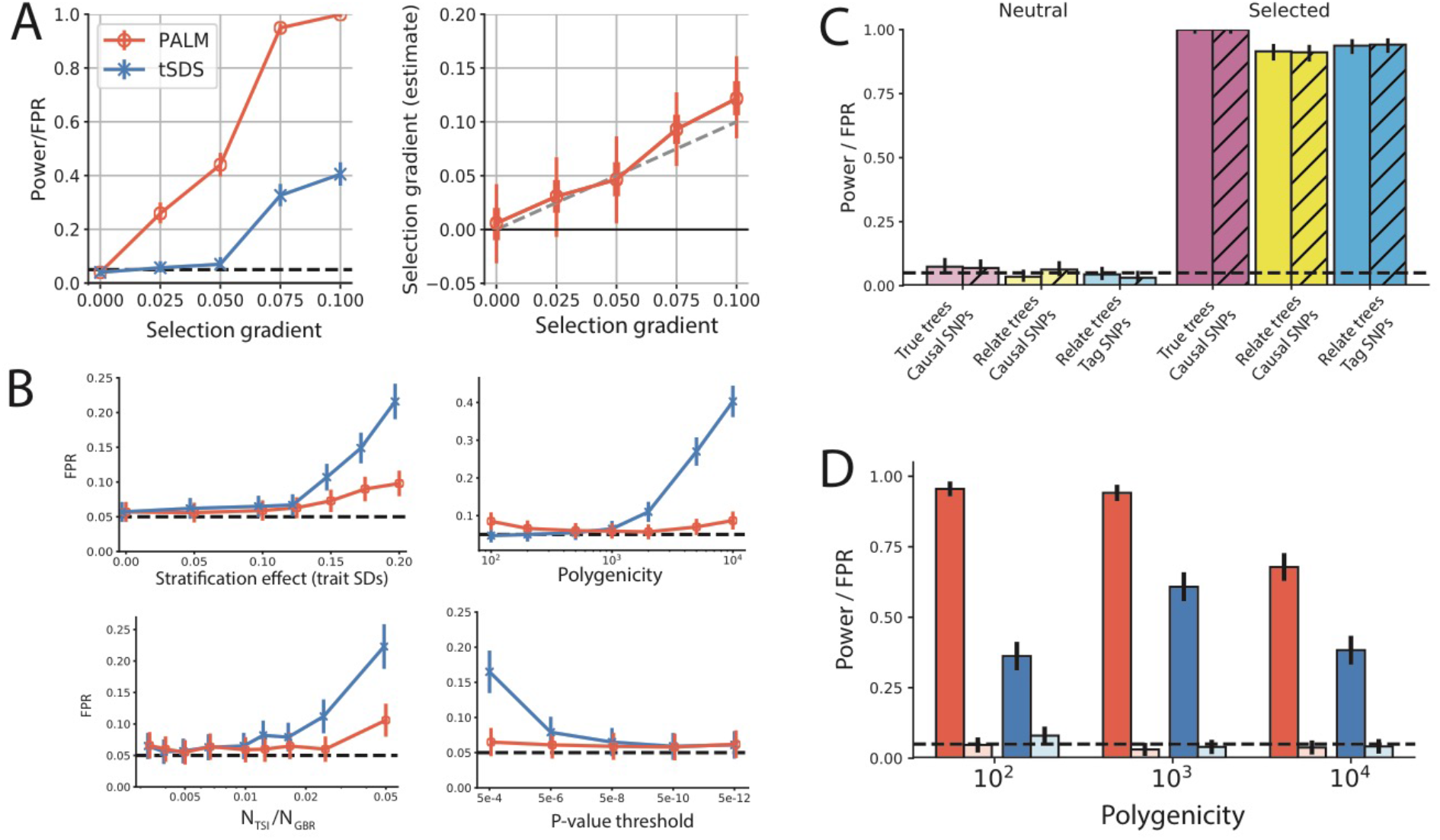
PALM power, calibration, and robustness to uncorrected stratification and ascertainment. (A) Left: Power/false positive rate (FPR) of PALM and tSDS. Error bars denote 95% Bonferroni-corrected confidence intervals. Right: PALM selection gradient estimates 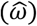. Error bars denote 25-75th percentiles (thick) and 5-95th percentiles (thin) of estimates; see Table 1 for more details of 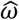 moments and error. Markers and colors in (A) also apply to (B,D). (B) False positive rate of PALM and tSDS applied to neutral simulations with uncorrected population stratification, simulated using 1000 Genomes data. We used baseline values of *σ_s_* = 0.1, *N_TSI_/N_GBR_* = 1%,*M* = 10^3^,*h*^2^ = 50%,using SNPs ascertained at *P* < 5 × 10^−8^. Error bars denote 95% Bonferroni-corrected confidence intervals. (C) Comparison of PALM using true *vs* Relate-inferred trees; causal *vs* GWAS-ascertained tag SNPs; and true marginal SNP effects (solid) *vs* GWAS-estimated SNP effects (hatched). Error bars denote 95% Bonferroni-corrected confidence intervals. (D) Varying polygenicity (*M*) of the polygenic trait. Error bars denote 95% Bonferroni-corrected confidence intervals. Baseline parameters for all simulations except (C) were our constant-size model with *M* = 10^3^, with Scz under positive selection and testing Scz for selection. In (A,B) we use Relate-inferred trees and estimated SNP effects at the causal SNPs; in D; we use Relate-inferred trees and estimated effects at tag SNPs. In all panels, we use a 5% nominal FPR (dashed horizontal line) and simulated 10^3^replicates.

We also examined the calibration and power of the marginal test in simulations of a polygenic trait with varying polygenicity (Fig. 1D). Across a wide range of polygenicities, PALM is well-powered to detect selection (>90% for 100 ≤ *M* ≤ 1000), with slightly lower power for extremely polygenic architectures (~ 65%for *M* = 10^4^) and the false positive rate (FPR) was well-calibrated in all circumstances (Fig. 1D). In comparisons to tSDS, we found substantially improved statistical power across this range of polygenicity values (Fig. 1D). We also conducted similar tests for a short pulse of selection (*ω* = 0.05for 35 generations, or ~1000 years assuming 29 years/generation) under a model of British demography^19^; we found that overall power was comparable to that of constant population size simulations with ω = 0.025, consistent with previous work showing that the product of selection strength and timespan largely determines statistical power (Supp. Fig 2).

#### Robustness to uncorrected GWAS stratification

We compared the power curve to the false positive rate (FPR) of both methods under a model of uncorrected GWAS stratification (Fig 1B). We simulated polygenic trait architectures and GWAS such that estimated SNP effects 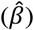 were both systematically biased and correlated with differences in the coalescence rate, stratified by DAF (e.g., SDS), matching the findings of^25,26^ that allele frequency differentiation between British (GBR) and Toscani in Italia (TSI) individuals was positively correlated with both 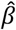 and SDS (Supp. Fig 1).

To model this scenario, we ascertained a set of 40,320 SNPs with MAF > 0.5% in the UKBB and SDS calculated by Field *et al*. (2016) using the UK10K cohort.^15^ We then sampled coalescence times at these SNPs in 1KG Phase 3 British (GBR) individuals using *Relate*. For each SNP, we simulated GWAS summary statistics by assuming that the GWAS cohort is comprised of some ratio, *N_TSI_/N_GBR_*, of TSI to GBR individuals, where population identity determines an individual’s stratified effect. This induces a correlation between SNP effects and the difference in allele frequency between TSI and GBR. Baseline parameter values were *σ_S_* = 0.1, *N_TSI_/N_GBR_* = 1%, *M* = 1,000, and *P* = 5 × 10^−8^. We varied the strength of the stratified effect (*σ_S_*, in phenotypic standard deviations) and found that both methods are well-calibrated when *σ_S_* is sufficiently small, but as *σ_S_* grows past 0.1 the FPR of tSDS was inflated over 100% more than that of PALM (Fig 1B).

We stress that this disparity is most likely not caused by higher sensitivity of tSDS, as we simulated polygenic adaptation under similar parameters and found PALM was better-powered to detect selection, with up to 8x improvement in power for smaller values of the selection gradient (Fig. 1A). We also found that for highly polygenic traits (e.g. *M* = 2 × 10^3^), the tSDS test is overconfident (>10% at 5% nominal), while PALM remains well-calibrated (Fig. 1B). We observe the same pattern as we increase the size of the cohort subgroup receiving the stratified effect (*N_TSI_/N_GBR_*); at *N_TSI_/N_GBR_* = 2.5% the tSDS test is overconfident (>10% at 5% nominal), while PALM remains well-calibrated (Fig. 1B).

Lastly, we tested the sensitivity of these methods to the stringency of the P-value threshold used, and found that the tSDS test was increasingly overconfident as the threshold was relaxed, whereas, PALM was well-calibrated regardless of P-value threshold (Fig. 1B).These results suggest that PALM is more robust to uncorrected stratification than the tSDS test, while obtaining superior statistical power even at lower sample sizes. However, we emphasize that PALM, like any other available test, is not fully robust to the effects of uncontrolled population stratification. Sufficiently strong uncorrected population stratification can lead to false inferences of polygenic selection when there is none.

#### Robustness to ascertainment bias and uncertainty in GWAS estimates

Next, we considered the effects of different levels of uncertainty and ascertainment on performance of PALM (Fig. 1C). We considered the effects of conditioning on the true local tree *vs* using Relate-inferred trees combined with importance sampling, conditioning on the true marginal SNP effect *vs* estimating this effect with noise in a GWAS; and conditioning on the causal SNP *vs* taking the top tag SNP in a local GWAS on linked SNPs. PALM was well-calibrated both using true trees and importance sampling, with highest statistical power (100%) using true trees and a slight drop in power under importance sampling (90-92%) (Fig 1C). Our test was well-calibrated despite bias (from Winner’s Curse) and noise in the estimated SNP effects, with no discernible difference from using the true SNP effects (Fig 1C); however, for smaller sample sizes (*N* << 10^5^) this may not be the case. Lastly, using the causal SNPs *vs* GWAS-ascertained tag SNPs did not diminish test power, and FPR remained well-calibrated (Fig 1C). We also explored the effects of GWAS sample size, which will affect the ascertainment process, and hence the degree of bias and uncertainty in ascertained SNP effect estimates (Supp. Tab. 2). We considered two different GWAS sizes; *N* = 10^4^ and 10^5^. We found that under lower sample size, the test was slightly inflated (e.g. empirical FPR of 3.1% (±1.4%) and 7.0% (±1.6%) at *N* = 10^5^, 10^4^ for Trait II respectively, where parentheses denote 95% CIs; Supp. Tab. 2). In terms of power, the test is still well-powered at lower sample sizes, but there is a noticeable drop (94.1% (±1.4%) and 69.0% (±3.0)% at at *N* = 10^5^, 10^4^ respectively; Supp. Tab. 2).

#### Robustness to model violations

We also conducted simulations of polygenic trait architectures under purifying selection, based on the model proposed by^7^ (Supp. Fig 3). Under such a scenario, an inverse relationship between effect size magnitude and derived allele frequency (DAF) is expected, in contrast to our baseline simulation model in which effect size is independent of frequency prior to the onset of selection. We found that across a range of polygenicities (M = 3 × 10^3^,10^4^,3 × 10^4^) and selection strengths (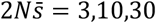, where 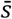 denotes mean selection coefficient of causal SNPs), PALM is not confounded by purifying selection and is well-calibrated to a nominal FPR of 5% (Supp. Fig 3); in fact, under very strong selection and/or low polygenicities, PALM is slightly conservative (Supp. Fig 3).

As our model and baseline simulations assume a single causal SNP per linked locus, we conducted simulations of allelic heterogeneity (Supp. Fig 4) using forward simulations in SLiM^31^. We simulated a trait architecture with *h*^2^ = 50%and a mutational target of 100 × 1Mbp linked loci, considering two cases: (1) 5% of incoming mutations are causal, and (2) 50% of incoming mutations are causal. In each of these scenarios we conducted simulations with neutral evolution and adaptation. We found that in each case, the test is well-calibrated under the null, and well-powered to detect selection (Supp. Fig 4).

Lastly, we explored the time specificity of PALM’s test for selection. Testing under a nominal model of selection in the last 50 generations, we consider the rate at which PALM’s estimate of selection timing can be biased by older selection (Supp. Fig. 5). We found that as selection recedes into the past, the FPR decays towards the nominal rate, with limited confounding when the pulse of selection occurred 200-250 generations ago.

#### Pleiotropy can cause bias in tests for polygenic adaptation

Traits with no fitness effect can undergo correlated response due to direct selection on pleiotropically related traits. Without accounting for pleiotropy, standard tests for polygenic adaptation cannot be interpreted as statements regarding direct selection. To illustrate how pleiotropy can affect tests for polygenic adaptation, we simulated pleiotropic trait architectures for 23 traits based on estimates of SNP heritability and genetic correlation for real human traits.^30^ This builds largely off our aforementioned simulation approach, with the introduction of a parameter *ϱ*, the degree of pleiotropy, i.e. the probability that a causal SNP is pleiotropic. As a brief illustration of how pleiotropy causes bias in polygenic selection estimates, we used our pleiotropic traits simulations to estimate maximum-likelihood selection coefficients for SNPs ascertained for associations to two genetically correlated traits, Trait I and II, modeled after schizophrenia and bipolar disorder (*r_g_* ≈ 80%; Supp. Fig. 6). We simulate a pulse of selection to increase Trait I (*ω* = 0.05, approximately +1 SD change in population mean over 50 generations, Supp. Tab. 1); Trait 2 has no causal effect on fitness. Selection coefficients were estimated by taking the maximum-likelihood estimate of s for each SNP independently, where the likelihood is estimated using our importance sampling approach. Here we show results for polygenicity *M* = 1000 and degree of pleiotropy *ϱ* = 60% (Supp. Fig 6).

Under the Lande approximation *s ≈ β*^⊤^*ω*, we expect a non-constant linear relationship between 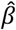 and 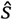 for traits under selection. But due to the strong correlation between these two traits, it is difficult to disentangle which of the traits has a causal effect on fitness (Supp. Fig 6A). We performed an ad-hoc test for a systematic relationship between 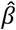 and 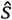 (Spearman test) to detect polygenic adaptation; while this test is well-powered to detect selection on Trait I, it is prone to spurious hits for selection on Trait II, which has no effect on fitness (Supp. Fig 6B). Thus, marginal tests for selection on traits can be significantly biased due to pleiotropy (in this case, genetic correlation).

#### Joint test for polygenic adaptation controls for pleiotropy

We also introduce a variant on our method, J-PALM, which is designed to disentangle correlated traits under selection and control for confounding due to pleiotropy. Briefly, J-PALM uses the same likelihood approach as PALM, but we jointly infer the selection gradient *ω*on a set of dtraits jointly, rather than inferring the selection gradient on a single trait marginally (see Model and Appendix for details). Under the joint model, the likelihood is still a function of the selection coefficient of each SNP, but we allow that these selection coefficients depend on the fitness effects of *d* traits jointly (see Model, Eq. 2).

We applied both our marginal test PALM and our joint test J-PALM to our cluster of four simulated traits, Traits I-IV, modeled after SNP heritabilities and genetic correlations for four psychiatric traits: schizophrenia, bipolar disorder, major depression and anorexia (Fig 2A). All traits have significantly positive genetic correlation to one another; here we highlight their genetic correlations to the selected trait, Trait I (Fig 2A; genetic correlations and SNP heritabilities directly from^29,30^). We assume a pulse of recent selection for increased Trait I prevalence, with all other traits selectively neutral. We tested traits marginally and jointly (i.e., all four simultaneously) for selection (Fig 2B,C). We found that marginal estimates are biased and cause inflation of the false positive rate (FPR) when testing for selection (Fig B,C). This bias largely follows the genetic correlation of the estimand trait to the selected trait (Fig 2A,B). Here we show results for polygenicity *M* = 1000 and degree of pleiotropy *ϱ* = 100% (Fig 2), but the results are similar for differing degrees of pleiotropy (holding *r_g_* constant), such as *ϱ* = 60% (Supp. Fig 7). This highlights that genetic correlation, regardless of the degree of pleiotropy, is the main cause of bias in marginal estimates of the selection gradient.

**Figure 2:**
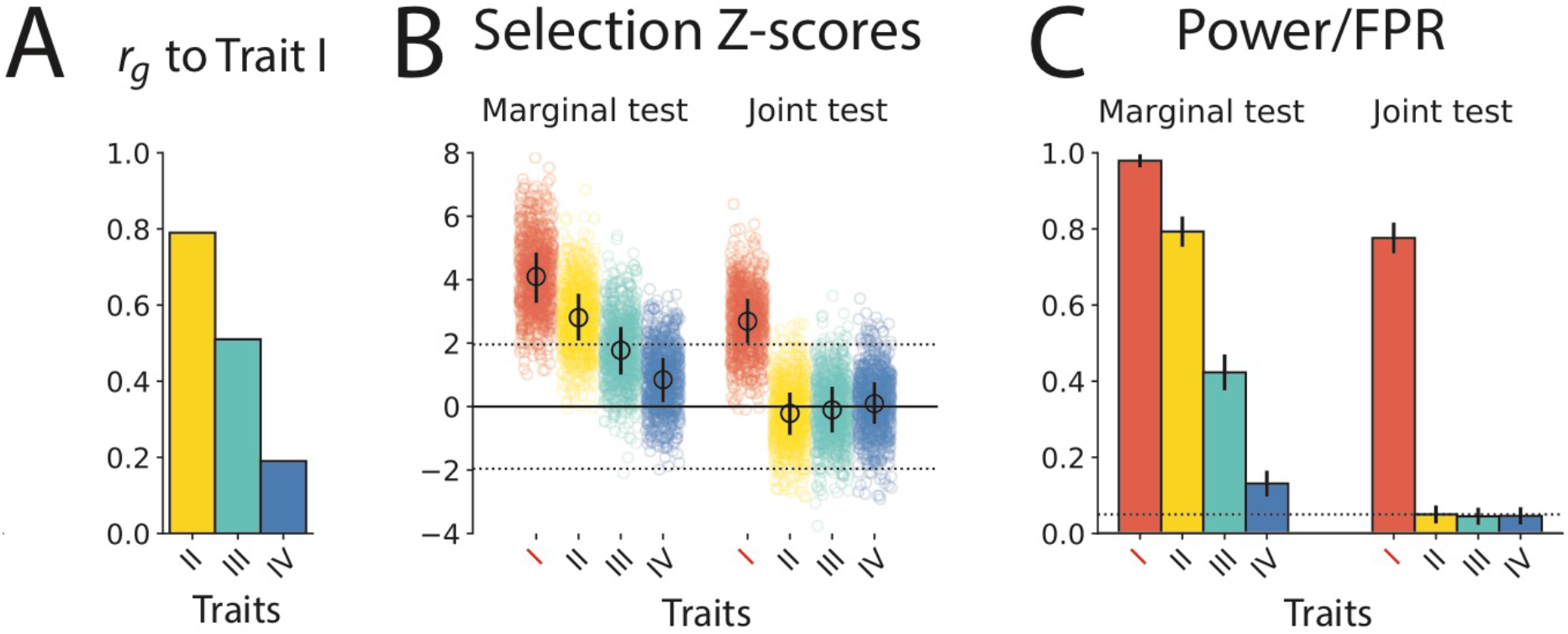
Joint testing for polygenic adaptation controls for pleiotropy. We simulated a cluster of four traits (I-IV) modeled after (A) real human heritability and genetic correlation estimates for schizophrenia (I), bipolar disorder (II), major depression (III), and anorexia (IV), with selection to increase Trait I in the last 50 generations. (B,C) We ran marginal and joint tests for selection on these four traits. While marginal selection tests were well-powered, they were strongly biased by even fairly low genetic correlations. (B,C) Conducting a joint test fully controls for genetic correlations while retaining high power to detect and isolate selection on Trait I. Simulations (1,000 replicates) were done under our constant effective population size model with *ϱ* = 60%, *M* = 1,000, with Trait I under positive selection.

Furthermore, our results show that if any trait in a genetically correlated cluster is under selection, marginal estimates of the selection gradient for the other traits is typically highly inflated. For example, a genetic correlation as low as *r_g_* = 19% is sufficient to inflate the FPR for a neutral trait by nearly 150% (Fig 2A,C). Most traits studied in GWAS have large genetic correlations; Watanabe, *et al*. (2019) found an average |*r_g_*| = 16% across 155,403 human trait pairs, with 15.5% of trait pairs significant (average |*r_g_*| = 38%).^32^ The extent of strong genetic correlation suggests that if any single heritable trait has evolved under selection, it is likely to cause substantial ripple effects in terms of bias of selection estimates on other heritable traits. By contrast, estimates of selection obtained via our joint test, fully correct for these biases, if the relevant selected trait is included in the analysis (Fig 2B,C). We applied the joint test to the same set of simulations and find it can reliably detect and attribute selection to Trait I (Fig 2B,C). The joint test preserved ~80% power even with the leading genetic correlate, Trait II, having *r_g_* = 79.4% to Trait I, and produces well-calibrated FPR regardless of *r_g_* (Fig 2C).

We explored performance of J-PALM under a wide array of simulation scenarios of different polygenic architectures and types of selection (Fig. 4), varying the degree of pleiotropy ρ (Fig 3A), *r_g_* to the selected trait (Fig 3B), polygenicity *M* (Fig 3C), and antagonistic selection (Fig 3D). Baseline values of parameters used were positive selection on Scz with other traits neutral, jointly testing Trait I and Trait III (*r_g_* = 51%), *ϱ* = 60%, and M=1,000. All of our pleiotropic simulations include an environmental noise correlation across traits of *p_e_* = 10%.Across this range of pleiotropic and polygenic architectures, we established that the joint test is well calibrated when no traits are under selection (Supp. Fig 8). Across different degrees of pleiotropy (40% ≤ *ϱ* ≤ 100%), we found J-PALM was well-calibrated and had good power to detect and attribute selection to Trait I (Fig 3A).

**Figure 3:**
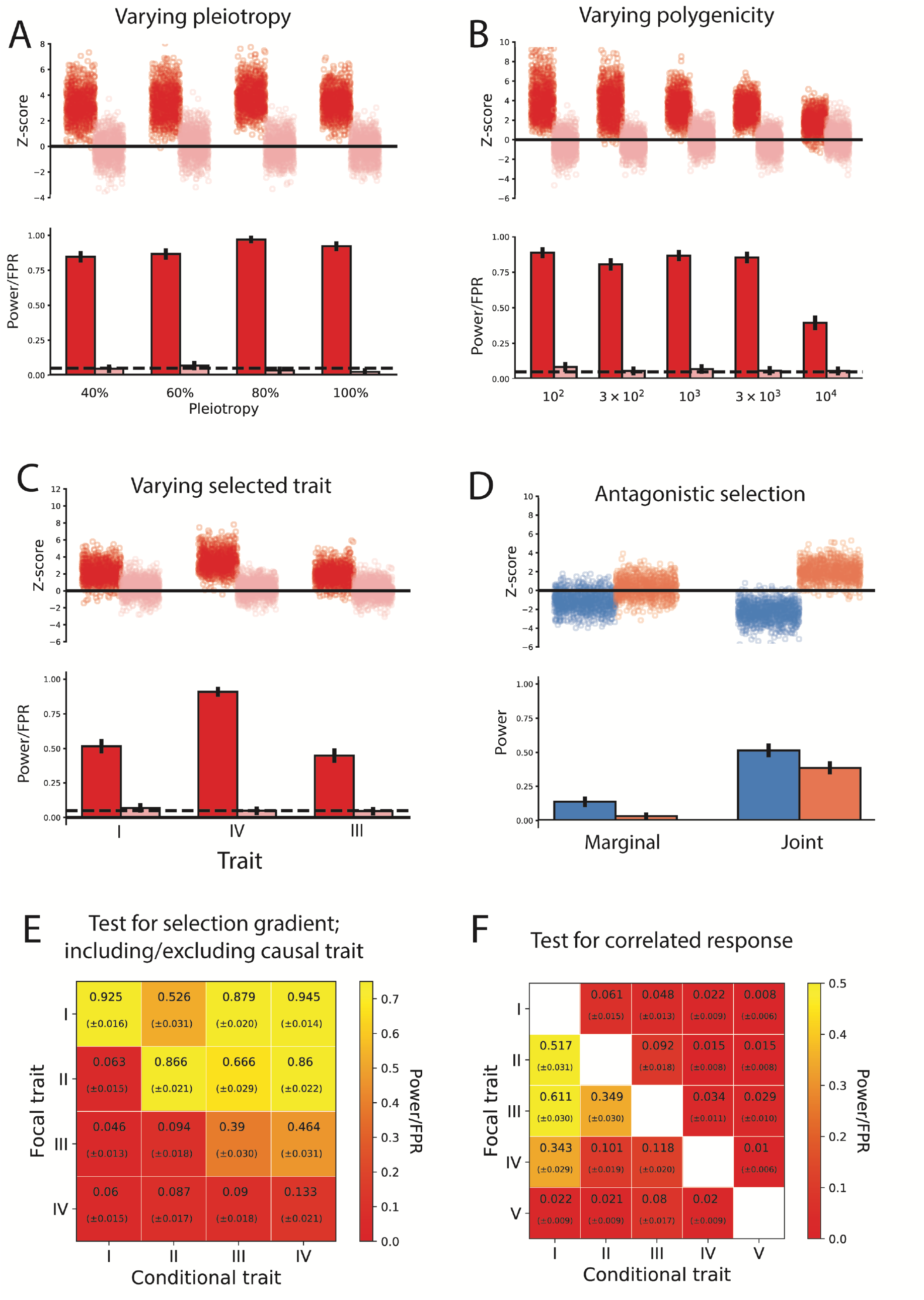
Simulations of joint testing power and calibration. (A) Differing the degree of pleiotropy *ϱ*, (B) the trait truly under selection, (C) the polygenicity *M* of the traits, (D) antagonistic selection on two traits with positive genetic correlation, (E) pairwise tests for selection (Trait I under selection), (F) pairwise tests for correlated response (Trait I under selection). (A-D) Red/pink/blue bars indicate estimates of selection for traits under positive selection/neutrality/negative selection, (E-F) Heatmap is colored by positive rate (also text in boxes; standard errors in parentheses). Dashed horizontal lines indicate 5% nominal significance level and black lines indicate 95% Bonferroni-corrected confidence intervals. Baseline parameters for all simulations (1,000 replicates under each scenario) were our constant-size model with *ϱ* = 60%, *M* = 1,000, with Trait I under positive selection. In panels (A,B) and (D) joint tests are performed on Trait I/Trait III and Trait I/Trait II, respectively. (E) Diagonal elements correspond to marginal test for selection.

Across a range of levels of polygenicity (100 ≤ *M* ≤ 10,000), PALM was well calibrated and had good power to detect and attribute selection to Trait I (>75% for *M* ≤ 3,000), although the power is somewhat attenuated for extremely polygenic architectures (~40% for *M* = 10,000) (Fig 3B). This pattern is also found in the marginal tests on the same data, and there is only a modest reduction in power when switching to the joint test (Fig 1C, Fig 3B). We note that the reduction in power is sensitive to the strength of genetic correlation; joint test of Trait I *vs* Trait II (*r_g_* = 79%) had greater reduction in power from the marginal test than that of Trait I *vs* Trait III (Fig 1C, Fig 3B,C, Supp. Fig 9). Our method fully corrects the biases suffered by marginal tests for polygenic adaptation, while retaining good power to detect adaptation even when genetic correlation is strong.

We also examined what happened when selection acted on different traits in the cluster, jointly testing each selected trait with Trait II (Fig 3C). The test is well-calibrated for all traits, but has less power to attribute selection to traits with a high genetic correlation to Trait II (e.g. Trait I, *h*^2^ = 45%, *r_g_* = 7%), or low heritability (e.g. Trait III, *h*^2^ = 17%, *r_g_* = 48%) (Fig 1E, Fig 3C). By contrast, traits with high heritability and/or low genetic correlation to Trait II (e.g. Trait IV, *h*^2^ = 49%, *r_g_* = 11%) have little loss in power in the joint test (Fig 1E, Fig 3C).

#### Detecting antagonistic selection

We also considered the possibility of antagonistic selection (i.e., selection to both increase Trait I and decrease Trait II, Fig. 3D). We hypothesized that marginal tests would be underpowered to detect this mode of selection acting on traits with strong genetic correlation, and that joint testing might uncover this signal. Indeed, power to detect selection in this regime is quite low using marginal testing, with 3-13% power at a 5% threshold (Fig 3D). However, the joint testing boosts power significantly, with 40-51% power at a 5% threshold (Fig 3D). We also tested the opposite scenario, where Trait I and Trait II are both under positive (complementary) selection; we found the joint test is well-powered to detect that multiple genetically correlated traits are under selection (Supp. Fig. 10). Thus, J-PALM provides several gains in power over the marginal test, such as uncovering antagonistic selection that is ‘cancelled out’ by genetic correlation, or confirming multiple traits are under selection.

#### Interpretation and limitations of the joint test

We also considered how our joint test performs when the causal trait (i.e., a trait with a causal effect on fitness) is excluded from the model. We conducted pairwise joint tests on each pair of Traits I-IV in simulations with Trait I under selection and all other traits neutral (Fig. 3E). Rows correspond to the trait for which the selection test is performed (the focal trait), and columns correspond to the other trait included in the joint model (the conditional trait). We also considered other scenarios, such as all traits neutral, complementary selection, and antagonistic selection (Supp. Fig. 11).

As we demonstrated previously, when the causal trait (Trait I) is included, the selection test is well-calibrated for neutral traits (Fig. 3E). However, we find that when Trait I is excluded, the selection test has high positive rates for traits that have no causal fitness effect, but are strongly genetically correlated with the causal trait (e.g. Trait II). In general, our results demonstrate that selection tends to be attributed to the trait with the strongest genetic correlation to the causal trait (e.g., Trait II); other traits with genetic correlation to the causal trait (e.g. Trait III) have some minor inflation of the positive rate, but selection is predominantly attributed to the closest proxy for the causal trait. These results highlight an important limitation of our model: Namely,the selection gradient estimates are not to be interpreted as causal fitness effects. As our simulated results show, this proposition is generally false when a trait with causal fitness effect and nonzero genetic correlation is excluded.

#### Testing for correlated response

Our method can also test for correlated response to selection, i.e., whether a trait has evolved (at least in part) due to selection on some other genetically correlated trait. We introduce the notion of an *effective selection gradient* (*ω_trait,model_*), which measures attributable amounts of selection to each trait included in a model. Consider two traits, A and B. Suppose Trait A is under selection and Trait B is neutral. If *r_g_* = 0, the effective selection gradient of B is 0, regardless of selection on Trait A or whether we include Trait A in the model, because no selection on A is attributable to B. Hence, *ω_B,marginaal_ = ω_B,joint_*. By contrast, if |*r_g_*| > 0, marginally Trait B has a nonzero effective selection gradient; however, in a joint model with Trait I, the effective selection gradient of Trait II is 0, since all direct selection can be attributed to Trait I. Hence, due to correlated response, there is a difference in the effective selection gradient in the two models: *ω_B,marginal_ ≠ ω_B,joint_*. However, the converse is not true for Trait I; both marginally and jointly with Trait II, all selection can be attributed to Trait I, and so *ω_A,marginal_ ≠ ω_A,joint_*. We developed a test statistic *R* (see Appendix) which tests for correlated response under the null hypothesis *H*_0_: *ω_j,marginal_ = ω_j,joint_*, i.e. that the marginal and joint effective selection gradients are equal.

We conducted tests of correlated response on each pair of traits I-V (we introduce Trait V, which has r_g_ = 0% to Trait I) (Fig. 3F). We found that the test for correlated response of Trait I is null, concordant with all other traits in the simulation being neutral (Fig. 3F). We also saw that Trait V, which has no genetic correlation to the directly selected trait, the test is null, concordant with the necessity of genetic correlation to drive correlated response (Fig. 3F). We saw that tests for correlated response generally grew in their power as r_g_ to Trait I increased. However, power is slightly lower for r_g_ = 80% than r_g_ = 50% (i.e., testing Trait II vs. Trait III for correlated response to Trait I) (Fig. 3F). This may indicate that for strongly genetically correlated traits, it is often ambiguous which one of the traits is evolving in correlated response. The test is also well-calibrated under neutral simulations (Supp. Fig. 12A), and well-powered to detect more complex forms of correlated response such as antagonistic and complementary selection (Supp. Fig. 12B,C). We also explored the performance of the correlated response test, along with the joint test for selection, in a K-way model with Traits I-IV tested jointly (Supp. Fig. 13). Our results indicate that our test statistic *R* can be used to detect whether a trait has been under correlated response; however, it is incorrect to make strongly causal interpretations of the test (e.g., “Trait III is under correlated response to Trait II”).

#### Effect of small or uneven GWAS sample size

We tested the effect of GWAS sample size on the joint test, considering not only lower sample size, but also uneven sample sizes (Supp. Tab. 2). Similar to the effect of lower sample size on the marginal test, we found that lower sample size for both traits reduced power and slightly inflated the FPR; e.g., testing for selection jointly on Trait I *vs* Trait II (simulating selection to increase Trait I), we found that at *N* = 10^4^ for Trait I and Trait II, the FPR for Trait II reached 8.0% (±1.8%) (Supp. Tab. 2). However, this was not always the case; e.g., for *N_I_* = 10^5^, *N_II_* = 10^4^, the FPR for Trait II was calibrated properly (4.6% ±1.4%) (Supp. Tab. 2).

Power to assign selection to the causal trait was reduced when that trait’s GWAS was underpowered; e.g., 51.6% (±1.6%) to 45.7% (±1.6%) when M,was dropped from 10^5^to 10^4^(*N_II_* = 10^5^) (Supp. Tab. 2). Interestingly, we found an even bigger drop in power associated with reduced sample size for the correlated trait (Trait II); when *N_II_*was reduced from 10^5^to 10^4^(M, = 10^4^), power to detect selection on Trait I dropped from 45.7% (±1.6%) to 27.7% (±1.4%) (Supp. Tab. 2). These results indicate that as long as sample size is reasonably large, estimates are well-calibrated; furthermore, by increasing sample size of GWAS for one trait, the joint test is able to leverage that towards improving power to detect selection on other traits that have overlapping genetic architecture.

### Empirical analysis of trait evolution in British ancestry

We analyzed 56 GWASs of metabolic, anthropometric, life history, behavioral, pigmentation- and immune response-related traits in humans (54 from the UKBB; see Supp. Tab. 3 for details) for signs of polygenic adaptation. We used GWAS summary statistics that were nominally corrected for population structure using either a linear mixed model^33^ or fixed PCs (K=20 PCs)^34^, and in some cases a family history-based approach^35^ to boost power for under-powered UKBB traits, such as type 2 diabetes. All traits used had at least 25 genome-wide significant (GWS) loci(*P* < 5 × 10^−8^) in independent LD blocks.^36^ For all of our empirical analyses, we used coalescent trees sampled using Relate for a sample of British ancestry (GBR, *n* = 89) from the 1000 Genomes Project, assuming pre-established estimates of GBR demographic history.^19–37^ We specifically tested for selection in the last 2000 years (i.e., 68.95 generations, assuming a generation time of 29 years). The selection gradient (*ω*) was estimated using maximum-likelihood, with standard errors estimated by block-bootstrapping. We first tested traits marginally for polygenic adaptation (Fig. 4). We include SNPs by pruning for LD using independent LD blocks, choosing the SNP with the lowest *p*-value in each independent block, and excluding blocks that do not have a SNP exceeding this threshold.^36^

**Figure 4:**
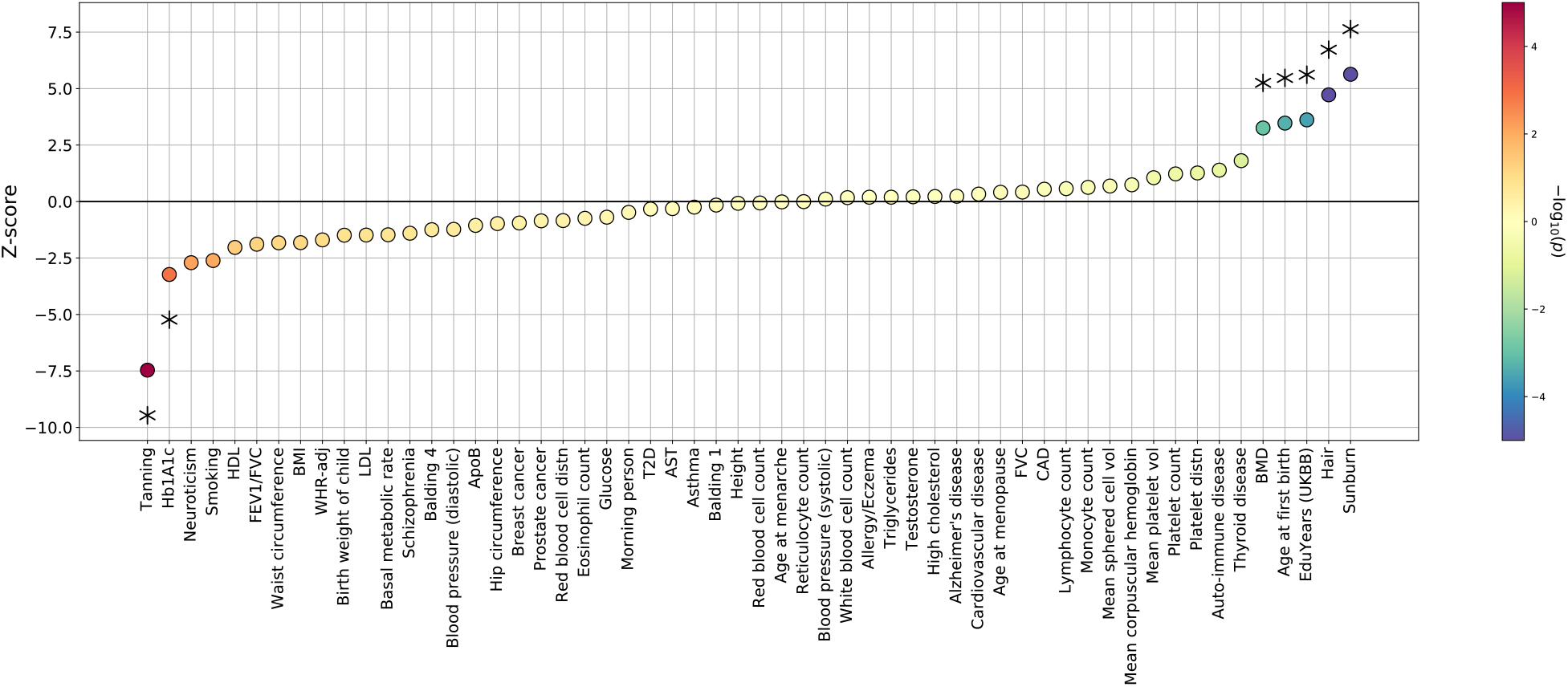
Estimates of the selection gradient on 56 human traits. The selection gradient 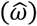 was estimated using 1000 Genomes Great British (GBR) individuals and summary statistics from various GWASs (see Supp. Tab. 4 for full results), with standard errors 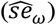 estimated via block-bootstrap 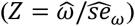. Starred traits indicate significance at FDR = 0.05.

#### Marginal tests for selection

We report our estimates of the selection gradient (Fig. 4) normalized by their standard errors, highlighting significant traits (FDR = 0.05) and other traits of interest, with results also presented in Supp. Tab. 4. In the marginal tests with PALM, we found strong signals of selection acting to decrease pigmentation (Fig. 4, Supp. Tab. 4). We reported traits with selection gradient p-value exceeding a multiple testing-corrected threshold (FDR = 0.05, Benjamini-Hochberg). Tanning showed the strongest signal of directional (in this case, negative) selection among all tested traits (*ω* = −0.357 (±0.046), *P* = 5.5 × 10^−15^; standard errors in parentheses). Sunburn (*ω* = +0.356 (±0.052), *P* = 1.1 × 10^−11^) and hair color (*ω* = +0.128 (±0.027), *P* = 2.2 × 10^−6^) also showed significant positive selection. Several life history traits also showed significant selection; e.g. age at first birth (*ω* = +0.0546 (±0.0149), *P* = 2.5 × 10^−4^) and EduYears (*ω* = +0.389 (±0.0107), *P* = 2.6 × 10^−4^). We also found significant selection acting on an anthropometric trait, bone mineral density heel-T Z-score (BMD, ω = +0.0728 (±0.0222), *P* = 1.1 × 10^−3^), and negative selection acting on glycated hemoglobin levels (HbA1c, ω = −0.0167 (±0.00518), *P* = 1.2 × 10^−3^) as well as neuroticism (*ω* = −0.0706 (±0.0254), *P* = 5.5 × 10^−3^).

Several traits of interest to have no or inconclusive evidence of directional selection. We found no evidence for any recent directional selection on height (*ω* = −0.00148 × 10^−3^ (±0.0190), *P* = 0.938). We also find inconclusive evidence for selection on body mass index (BMI, (*ω* = −0.0585 (±0.0331), *P* = 0.077), in contrast to previous findings that BMI has been under significant selection to decrease.^16^

#### Joint tests for selection

We analyzed 137 trait pairs (Bonferroni *P_rg_* < 0.005 and |*r_g_*| > 0.2)^32^ using J-PALM to examine if marginal signals of selection were due to a correlated response to selection on another trait (Table 2, Supp. Tab. 5). To aid clarity, we introduce the notion of focal *vs* conditional traits in a joint test. For example, if we estimate the selection gradient of Trait 1 and Trait 2, (*ω*_1_, *ω*_2_), then *ω*_1_is the estimate for Trait 1 (the focal trait), jointly tested estimated with Trait 2 (the conditional trait); similarly *ω*_2_is the estimate for Trait 2 (the focal trait), jointly tested estimated with Trait 1 (the conditional trait). We establish significance of correlated response using a Wald test on the statistic P. the difference in the joint and marginal selection estimates for a focal trait, where the joint analysis is performed with some other conditional trait (see “Testing for correlated response” and Appendix for more details). Selected results are presented in Table 2, and results for the full analysis of all 137 trait pairs are available in Supp. Tab. 5.

**Table 2:**
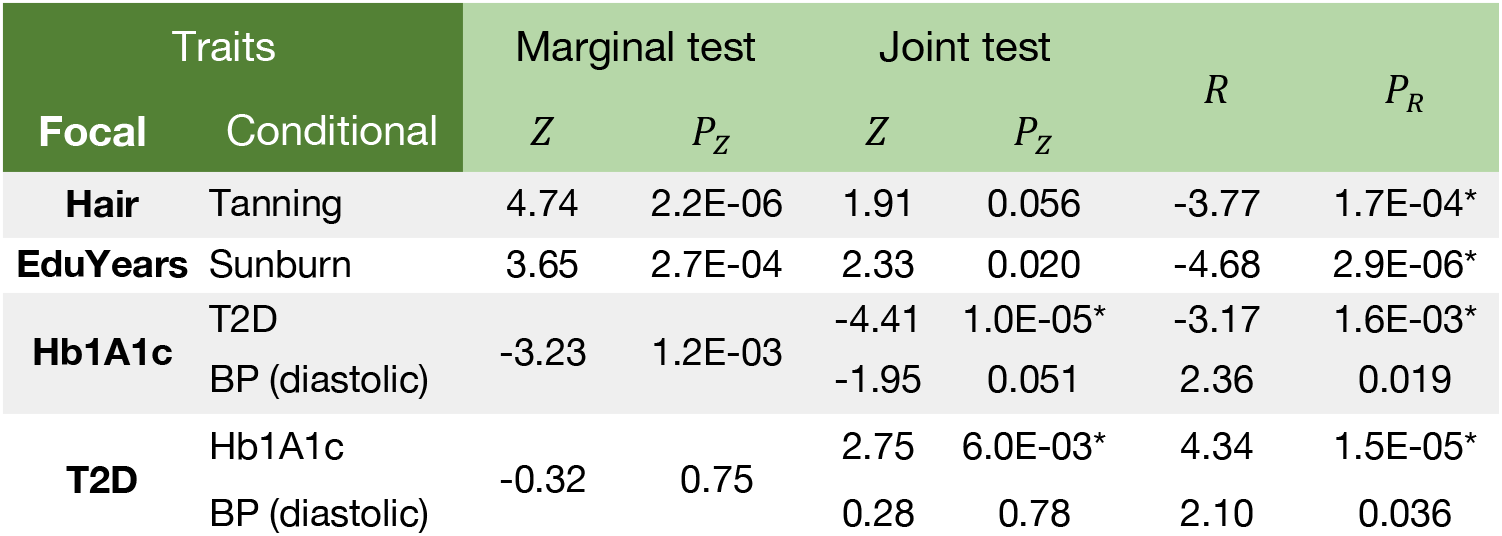
Selected trait pairs under correlated response in Great British ancestry. Selection on the focal trait is estimated jointly with the conditional trait. We report the Z-scores under both the marginal and joint tests, as well as the *R* statistic of the difference in joint *vs* marginal selection gradient estimates, and their *P*-values. Results for all trait pairs are available in Supp. Tab. 5. T2D = Type 2 diabetes, HbA1c = glycated hemoglobin, BP = blood pressure. Asterisk (*) denotes significance at FDR = 0.05 (*n* = 2 × 137 = 274tests on 137 trait pairs with Bonferroni-significant 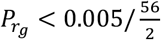 and |*r_g_* > 0.20|.

We found several significant signals (FDR = 0.05) of correlated response (Table 2, full results in Supp. Tab. 5). For example, although EduYears had strong evidence for selection in the marginal test (*P_marginal_* = 2.6 × 10^−4^), we found after conditioning on sunburn ability (*r_g_* = 0.24, *P* = 2.3 × 10^−4^)^32^ a significant attenuation of this estimate (*P_joint_* = 0.020, *P_R_* = 2.6 × 10^−6^). These results suggest that a large part of the signal of selection on EduYears is likely due to indirect selection via correlated response, *vs* direct selection. However, we stress that these results do not provide evidence of direct selection on the conditional trait, here e.g. childhood sunburn occasions (sunburn) (see e.g. Fig. 3E).

We also find significant attenuation of selection signals for pigmentation traits in our joint analyses (Table 2). In our joint analysis of hair color and tanning (*r_g_* = −0.17, *P* = 3.6 × 10^−3^)^32^, we found that after conditioning on tanning, there is no residual evidence for direct selection on hair color (*P_marginal_* = 2.2 × 10^−6^; *P_joint_* = 0.056; *P_R_* = 1.7 × 10^−4^). (The same caveat above regarding the interpretation of correlated response applies here to tanning ability).

We identified one case in which the joint analysis uncovers selection acting on a trait that did not show significant selection marginally; we found that type 2 diabetes (T2D), conditioning on HbA1c (*r_g_* = 0.69)^38^, shows significant selection to increase in prevalence(*P_marginal_* = 0.75; *P_joint_* = 0.0060; *P_R_* = 1.5 × 10^−5^; see Table 2). Estimates of negative selection on HbA1c are also enhanced after accounting for T2D (*P_marginal_* = 1.2 × 10^−3^; *P_joint_* = 1.0 × 10^−5^; *P_R_* = 0.0016; see Table 2). This ‘cancelling-out’ effect of opposing selection on T2D and HbA1c, two traits with strong positive genetic correlation, is the second-strongest signal of correlated response in our joint analyses.

We also illustrate our estimates of selection coefficients for ascertained T2D/HbA1c SNPs, found independently of one another, and their fit to our inferred model of antagonistic selection on T2D/HbA1c (Fig. 5A). In general, T2D-increasing and/or HbA1c-decreasing SNPs are under positive selection, and vise versa; however, a subset of HbA1c-increasing SNPs show extremely strong signs of positive selection (*s* > 0.03); these SNPs tend to have visibly higher positive effects on T2D than other SNPs with comparable HbA1c effect. In a joint analysis of HbA1c and diastolic blood pressure (as a proxy for hypertension), our estimate of direct selection on HbA1c was significantly attenuated at a nominal level (*P* = 0.019, Table 2), although it did not meet our FDR cutoff. We also did a joint analysis of T2D and diastolic blood pressure, finding a significant boost in the estimate of direct selection on T2D (*P* = 0.036, Table 2), although it did not meet our FDR cutoff.

**Figure 5:**
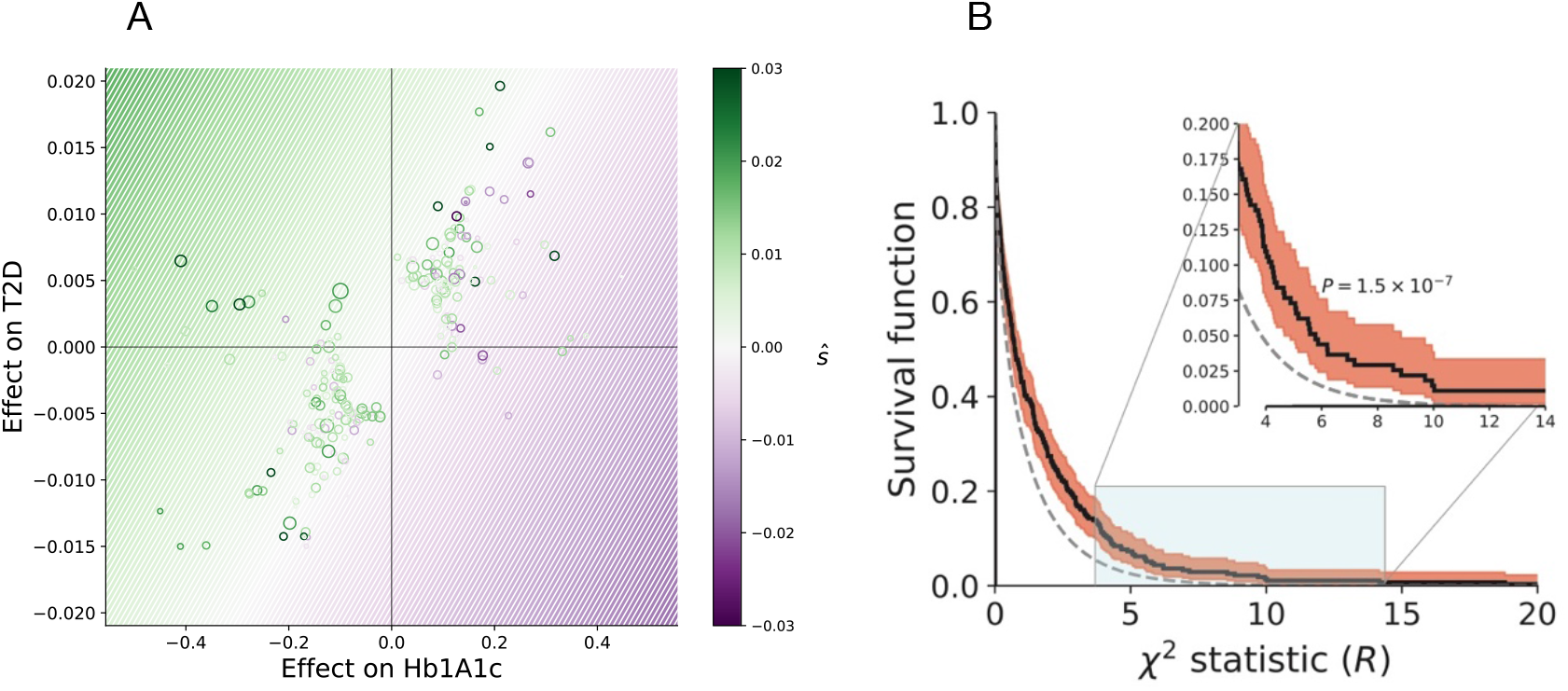
Correlated response in real traits. **(A)** Expanded view of antagonistic selection on glycated hemoglobin (HbA1c) *vs* type 2 diabetes (T2D). We estimate individual SNP selection coefficients by taking the maximum-likelihood estimate *ŝ* for each SNP. We plot this value against the joint SNP effect estimates for HbA1c and T2D. Colored lines represent isocontours of 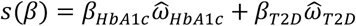, the estimate of the Lande transformation from SNP effects to selection coefficients, where 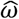is inferred jointly for the two traits (Table 2). **(B)** Enrichment of correlated response in analysis of genetically-correlated traits. Enrichment in the tails of the distribution of our test statistic for correlated response *R*(*R* = 1.5 × 10^−7^, binomial test) which had 2.6-fold enrichment at the nominal 5% level. We assessed *n* = 2×137 = 274estimates of correlated response on 137 trait pairs with Bonferroni-significant 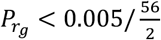 and |*r_g_* > 0.20|. Red area indicates pointwise 95% CI of the survival curve.

Lastly, we tested our set of P statistics among the pairs of genetically correlated traits for enrichment in the tail over the null (Fig. 5B). At the nominal 5% FPR level, we found significant (2.6-fold) enrichment for correlated response acting on these traits (*P* = 1.5 × 10^−7^, one-sided binomial test), suggesting that many additional traits in this analysis have evolved under indirect selection due to correlated response.

## Discussion

We have presented a method, PALM, for estimating the directional selection gradient acting on a polygenic trait. Our method works by estimating likelihood functions for the selection coefficients of a set of GWAS SNPs, and then aggregating these functions along with GWAS-estimated SNP effects to find the likelihood of the selection gradient. Through simulations, we showed that PALM offers improved power over current methods across a range of selection gradients(*ω* = 0.025 − 0.10) and polygenicities (*M* = 10^2^ − 10^4^), and is the first method to our knowledge that can estimate *ω*from nucleotide data. We conducted robustness checks and showed that PALM is robust to typical sources of uncertainty and bias in GWAS summary statistics (e.g. sampling variation, ascertainment bias/Winner’s Curse) allelic heterogeneity, purifying selection, and underpowered GWAS.

We also introduced a method, J-PALM, to jointly estimate the selection gradient on multiple traits in order to control for pleiotropy. We showed that, across a wide range of polygenic architectures (*M* = 10^2^ − 10^4^, *ϱ* = 40% − 100%), J-PALM can reliably detect and assign selection to the causal trait when it is considered in the analysis, and can be used to uncover genetically correlated traits under antagonistic selection where the marginal approach (e.g. PALM) is underpowered. We considered several additional sources of bias unique to multi-trait analyses (i.e. uneven GWAS sample sizes, correlation in trait environmental noise) and found J-PALM robust to these as well.

We note several areas in which the study of polygenic adaptation can be advanced. Our operative model of polygenic adaptation is based on the Lande approximation, which over long time-courses will overestimate the efficiency of adaptation under stabilizing selection with a shift in the optimum.^12–39^ A model that incorporates these dynamics will potentially be better suited to detecting polygenic adaptation over longer time-courses, such as analyses of ancient DNA samples. Furthermore, under stabilizing selection more SNP heritability is expected to be sequestered to low-frequency alleles, and so common SNPs are expected to change less under adaptation than in our simulation model.^5,12^

Advances might also be made through more nuanced models that make fuller use of GWAS summary statistics and LD among GWAS marker. We showed our thresholding and pruning scheme for selecting sites did not substantially decrease our method’s power. Pre-existing methods for fine-mapping or ascertaining pleiotropic loci might increase power even further.^40^ It is also possible that for traits with extremely high polygenicity and/or low heritability, it will be necessary to utilize summary statistics that are sub-significant, and account for uncertainty in the location of the causal site.

We showed that PALM is substantially less prone to bias due to uncorrected GWAS stratification than comparable methods such as tSDS. However, we stress that PALM can nonetheless be biased under sufficiently strong uncorrected stratification. Other forms of stratification that we did not explore, such as gene-by-environment (GxE) interactions, may be more difficult to account for via standard kinship-based approaches; however, new methods have recently arisen to this particular end.^41^

Another limitation of our model is the interpretation of the estimates of the selection gradient and correlated response. We showed through simulations that when a genetically correlated trait with causal fitness effect is excluded from the analysis, estimates of direct selection have no causal interpretation. To address this, we introduced the notion of an effective selection gradient, which depends on which traits are modeled together. Estimates of the effective selection gradient allow us to determine whether a focal trait has evolved under correlated response another trait; however, this does not have the causal interpretation that the focal trait is under correlation response to a particular conditional trait.

Applying PALM to study evolution of 56 human traits in British ancestry, we found 8 traits under significant directional selection, recovering several previously-reported targets, such as pigmentation traits, educational attainment, and glycated hemoglobin (HbA1c), in agreement with previous findings of selection on these traits in Europe.^15,16,42^ We also report several novel targets of directional selection, such as increased bone mineral density and decreased neuroticism. Despite historical claims of selection to increase height in Europe^22^, we found no evidence for selection to increase height, consistent with recent analyses which showed that signals of directional selection on height have been drastically inflated by uncorrected population structure in GWAS summary statistics.^25,26^

We applied our joint test J-PALM to study 137 pairs of genetically correlated traits for signatures of correlated response. We found a highly significant enrichment of correlated response acting on these traits. Particularly, we found significant correlated response acting on pigmentation and life history traits (hair color, educational attainment). We showed that signal of selection on traits such as hair color and educational attainment, which have been widely reported to date^15,16,42,43^, are due in significant part to correlated response to selection on other traits, *vs* direct selection acting on these traits.

One proposed theory for the diversification and increase of blonde hair color in Europe is sexual selection.^44,45^ However, our results do not support this, as we show that evidence for selection on hair color is attributable mostly to correlated response, beyond which there is little evidence for direct selection on this trait. This echos previous analysis showing selection at individual hair color loci may be indirect, via their pleiotropic effects (e.g. blonde hair gene *KITLG* responding to selection for tolerance to climate and UV radiation^46^), and conflict with arguments that hair color has been under direct sexual selection.

In our marginal test for selection, we detected significant selection to increase educational attainment, consistent with some previous work.^16^ However, in a joint test with sunburn (i.e., “childhood sunburn occasions,” the number of times the individual was sunburned as a child), strong signals of selection to increase educational attainment were significantly obviated. We conclude that signals of selection on educational attainment are driven significantly by correlated response. We caution that “childhood sunburn occasions” is a survey question, and is likely affected by many exogeneous factors beyond skin pigmentation (e.g., opportunity to visit the beach or use sunscreen). We propose that gene-by-environment (GxE) interactions may be driving these signals of correlated response. Lewontin (1970), responding to Jensen (1968), pointed out that then-current estimates of intelligence quotient (IQ) heritability were inflated by GxE.^47,48^ Indeed, in modern-day GWAS, we see that educational attainment polygenic scores in the UKBB are only 50% as predictive in adoptees as in non-adoptees, indicating a significant role of GxE in the expression of educational attainment, as well as estimates of its heritability and genetic correlations^49^ Hence, genetic correlation of sunburn and educational attainment may be overestimated (e.g., 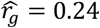 using UKBB GWAS^32^). We do not have data to elucidate the mechanism of this proposed GxE interaction, but hypothesize that educational opportunities and other environmental influences could be affected by skin pigmentation. Even in the absence of GxE, we stress that our results are not interpretable as evidence of direct selection on “childhood sunburn occasions”--let alone skin pigmentation--following from our simulation study. Lastly, the inferred correlation between the traits and/or the signals of selection could be affected by uncorrected GWAS stratification.^25–26^

We found one case of significant antagonistic selection: T2D shows significant selection to increase, but this signal was initially occluded by the positive genetic correlation of T2D with negatively-selected glycated hemoglobin (HbA1c). Our joint analysis with J-PALM disentangles this antagonism between T2D and HbA1c, revealing latent adaptation of T2D. T2D is a complex disease with a complex etiology, involving obesity and various metabolic risk factors. Selection may have favored some of these factors under previous environmental conditions where both obesity and diets rich in simple sugars were uncommon (also known as the thrifty gene hypothesis).^50^ HbA1c is a biomarker commonly used to not only diagnose pre-diabetes/diabetes, but also to monitor chronic hyperglycemia as a risk factor for vascular damage.^51^ T2D and HbA1c are strongly, although imperfectly genetically correlated (*r_g_* = 69%), and HbA1c is associated with hypertension and other cardiovascular disease independently of T2D incidence.^38^ It is therefore possible that selection might have favored some of the traits underlying increased T2D risk, but acted against some of the more specific negative effects of T2D which now are measured by HbA1c.^38,51,52^ These results provide evidence in support of the thrifty gene hypothesis.^50^

## Methods

### Simulations

#### Pleiotropic polygenic trait architecture

We sample effect sizes jointly for *d* = 23 polygenic traits with previously estimated SNP heritability and genetic correlations.^29–30^ We consider different values of polygenicity (*M*, the number of causal SNPs) and degrees of pleiotropy (*ϱ*, the probability that a causal SNP is pleiotropic). Let *G* be the additive genetic covariance matrix (diagonal entries are the SNP heritabilities for each trait). Then the genetic correlation of traits *i, j* is 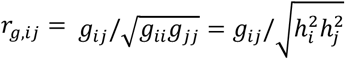. Under our simulation model, we assume that if a SNP is pleiotropic, then *β* ~ MPA(0, G^*^/(*Mv*)), where *g^*^_ii_ = g_ii_* · (1 − (1 − *ϱ*)|*d*)|*ϱ, g^*^_i=j_ = g^*^_i≠j_/ϱ*. If a SNP is non-pleiotropic and is causal for trait *j*, then 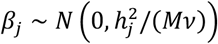 where 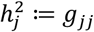, and *β_−j_*. = 0. We assume that if a SNP is non-pleiotropic, it is causal for a particular trait 7’with uniform probability 1/*d*. Under this model, we can see that averaging over pleiotropic and non-pleiotropic loci, we recover the overall genetic covariance *G*:

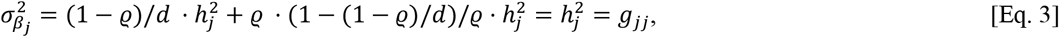

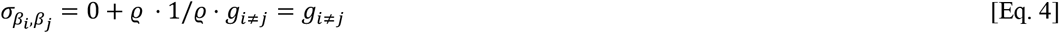

Note that here *β* is standardized by the phenotypic variance, but not the genotypic variance. Thus we normalize the variance by a factor of *v* = 2 · *E*[*pq*], assuming some stationary distribution for *p*. We assume the stationary distribution *f*(*p*) ∝ 1/*p*, which yields *v* = 4 *log N_e_*, where *N_e_* is the diploid effective population size. This choice of *v* ensures 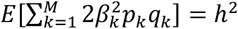 under the nominal allele frequency spectrum. The equation holds because we assume independence of effects and allele frequencies; we also performed simulations where *β* and *p* are allowed to depend strongly on each other due to purifying selection.

#### Simulation of confounding due to population structure and uncorrected GWAS stratification

Previous estimates of selection to increase height in Europe have been biased by a combination of uncorrected stratification and GWAS and systematic differences in the coalescent rate at SNPs that depended on their allele frequency difference in 1000 Genomes (1KG) British (GBR) vs. Southern Italy (TSI) populations.^25,26^ We developed a simulation model based on empirical data from the 1KG data in order to assess the robustness of our method compared to tSDS-based tests for polygenic selection.^15^ We model uncorrected stratification in summary statistics for a simulated polygenic trait architecture by drawing random SNP effects

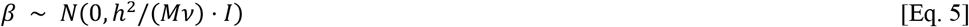

where *I*is the identity matrix. We assume that the phenotype follows the form

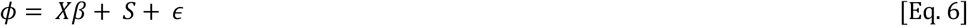

where *S* is some environmentally determined stratified effect experienced by an individual based on whether they belong to a subpopulation. If *N*_1_, *N*_2_ individuals (*N*_1_, + *N*_2_ = *N*) belong to subpopulations 1 and 2 (e.g., GBR and TSI) respectively, then 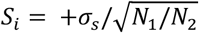 if 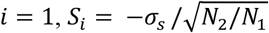 if *i* = 2. (It can be shown then that phenotypic mean remains 0, and variance due to stratification is 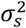.) Under this form of stratification, assuming random mating of genotypes, the expected effect estimate is biased:

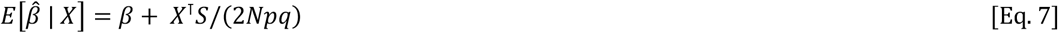

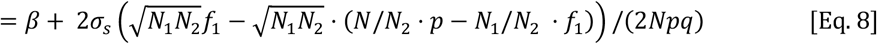

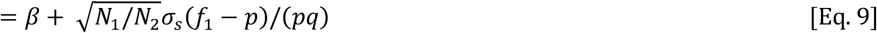

where *p* = 1 − *q* = (*N*_1_, *f*_1_, + *N*_2_*f*_2_)/*N*) is the overall frequency of the SNP, and *f*_1_ is the frequency of the SNP in subpopulation 1. The nominal standard error of 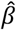 is the usual 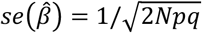.

Hence, we can simulate GWAS-estimated SNP effects with uncorrected stratification using

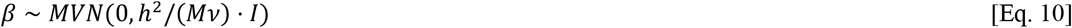

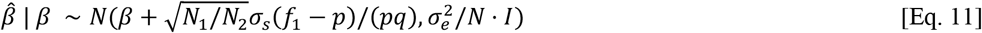

where 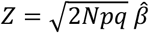 and *σ_e_*^2^: = 1 − *h*^2^ − *σ_s_*^2^. Although in this simple model of GWAS with uncorrected stratification, we assume no LD between causal sites, the bias in the effect estimates does not depend on LD. We note that this is equivalent to the model of Bulik-Sullivan, *et al*. (2015)^29^, generalized to uneven sample sizes from subpopulations.

#### Population genetic model of selection and ascertainment bias via GWAS

Given μ, we simulate selection following the multivariate Lande approximation (see Model). Because we simulate polygenic architectures of *M* ≥ 100 without linked causal loci, out assumption of infinitesimal genetic architecture is appropriate. (We also explore the performance of our model when we allow LD between causal SNPs; see Supp. Fig. 4). We then simulate the trajectory of the allele forward in time using a normal approximation to the Wright-Fisher model with selection, i.e. *p*_*t*+1_ ~ *N*(*p_t_ + sp_t_*(1 − *p_t_*), *p_t_*(1 − *p_t_*)/4*N_e_*), where s is calculated using the multivariate Lande approximation. For most of our simulations, we simulate forward for 50 generations (i.e., we assume selection began 50 generations before the present), unless otherwise stated. Let *p* be the present-day allele frequency. We simulate the ascertainment of this SNP in a GWAS by simulating the SNP Z-scores 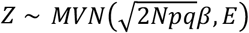, where *E_ii_* = 1, *E_i≠j_* = *ρ_e_*, where *ρ_e_* is a term that allows for cross-trait correlations in environmental noise. (Note that here *Z*is the usual Z-score of 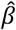, not to be confused with the selection Z-score we present earlier.) Unless stated otherwise, we set *N* = 10^5^, *ρ_e_* = 0.1 in all simulations. We use a p-value threshold of 5 × 10^−8^ to ascertain a SNP; this must be surpassed by at least one trait. If a SNP is ascertained, we simulate its trajectory backwards in time using the normal approximation to the neutral Wright-Fisher diffusion conditional on loss, *p*_*t*−1_ ~ *N*(*p_t_*(1 − 1/4*N_e_*), *p_t_*(1 − *p_t_*)/4*N_e_*). We use the coalescent simulator *mssel* to simulate a sample of haplotypes conditional on this allele frequency trajectory.^20^ We use *n* = 400 haplotypes and *μ* = *r* = 10^−8^/bp/gen and simulate regions of 1Mbp, centered on the causal SNP at the position 5 × 10^5^.

To simulate ascertainment of non-causal SNPs in a GWAS, we take the trait with the top Z-score at the causal SNP and jointly simulate Z-scores for that trait for all linked SNPs within a 200kbp window centered on the causal SNP and surpassing a MAF threshold (MAF ≥ 0.01). We ascertain the SNP with the top Z-score (sometimes the causal SNP), and then simulate the Z-scores for all traits, conditioned on the Z-score for the one aforementioned trait. We simulate this way rather than jointly simulating Z-scores for all traits at all SNPs because for two reasons; the top SNP will typically have the same top trait association as the causal, and jointly simulating all trait-by-SNP Z-scores increases computational time by >400 for the parameters we used.

To further reduce computational burden, we simulated libraries of 10 × *M* causal loci and resampled sets of Mloci without replacement (some proportion of which meet the ascertainment criteria), in order to model sampling variation in the test statistics.

#### Inference of local genealogies

Given a set of simulated haplotypes, we use the software package *Relate*^19^ to infer local genealogies along the sequence. Using positions of the SNPs ascertained through GWAS, we use the add-on module *SampleBranchLengths* to draw *m* = 5,000 MCMC samples of the branch lengths of the local tree at the ascertained sites. We then extract coalescence times from these MCMC samples (thinned down to *m* = *500* approximately independent samples), and partition the coalescence times for each sample tree based on whether they occur between lineages subtending the derived/ancestral alleles. We note that *Relate*, unlike *ARGweaver*, does not sample over different ARG or tree topologies, and it samples branch lengths for two distinct local trees independently, conditional on the observed data.

#### Comparisons to tSDS in simulations

In order to calculate tSDS values for our simulated polygenic traits, we computed the Gamma shape parameters for a model with constant *N_e_* = 10^4^ using 250 simulations at a range of DAFs from 1% to 99%, with 2% steps between frequencies, and a sample size of *n* = 400 haplotypes. We randomly paired haplotypes in the sample to form diploid individuals and found singletons carried by each diploid. We then calculate raw SDS using the *compute_SDS.R* script with our custom Gamma-shapes file. To calculate SDS we find the Z-score of a SNP’s raw SDS value, where the mean and standard deviation are estimated from an aggregated set of 29,478 completely unlinked SNPs from our neutral trait simulations. To calculate tSDS we calculate the P-value of the Spearman correlation of (sign(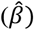),SDS).

## Acknowledgements

We thank Doc Edge for feedback on the manuscript, and Jeremy Berg, Jennifer Blanc, Yun Deng, Daniel Geschwind, Iain Mathieson, Priya Moorjani, Monty Slatkin, and Lawrence Uricchio for helpful discussions.

## Author contributions

AJS, NAZ, and RN conceptualized the study; AJS, NAZ, and RN designed methodology; AJS performed experiments and analysis; AJS developed the software; AJS and LS curated the data; NAZ and RN supervised the study; AJS wrote the manuscript; AJS, LS, NAZ, and RN edited the manuscript.

## Resources & URLs

- Open-source code and documentation for PALM/J-PALM is available at http://www.github.com/35ajstern/palm.
- Formatted summary statistics/metadata and 1000 Genomes GBR selection likelihoods for ascertained SNPs are available for download on DataDryad: https://doi.org/10.6078/D11M62.
- Other web resources: 1000 Genomes Phase 3 data, ftp://ftp.1000genomes.ebi.ac.uk/vol1/ftp/phase3/; Neale Lab GWAS Round 2, https://tinyurl.com/ycg5bxq5; BOLT-LMM summary statistics, https://data.broadinstitute.org/alkesgroup/UKBB/UKBB_409K/; LT-FH summary statistics, https://data.broadinstitute.org/alkesgroup/UKBB/LTFH/sumstats/; Alzheimer’s Disease GWAS summary statistics, https://ctg.cncr.nl/software/summary_statistics; PGC summary statistics, https://www.med.unc.edu/pgc/download-results/; GWAS Atlas, http://atlas.ctglab.nl/; Relate software, https://myersgroup.github.io/relate/; SDS scripts, https://github.com/yairf/SDS.

