## Supplementary material for "Disentangling selection on genetically correlated polygenic traits using whole-genome genealogies": SUPP_text_figs_tables: supp.pdf

### Table of Contents


Appendix 3

Inference 3

Importance sampling estimation of the likelihood function of selection 3

Accounting for multiple SNPs in LD 4

Selection gradient and correlated selection standard errors 5

Coalescent likelihood models 5

*Relate* prior 5

Coalescent selection likelihood under deterministic model 6

Supplementary Figures 8

Supp. Fig. 1: Distribution of frequencies and SDS in 1000 Genomes SNP set. 8

Supp. Fig. 2: Calibration and power under GBR demography. 9

Supp. Fig. 3: Robustness to purifying selection. 10

Supp. Fig. 4: Calibration and power under allelic heterogeneity. 11

Supp. Fig. 5: Time specificity of test for recent selection. 12

Supp. Fig. 6: Pleiotropy causes bias in tests for polygenic adaptation. 13

Supp. Fig. 7: Marginal vs. joint test comparison, lower pleiotropy. 14

Supp. Fig. 8: Calibration of joint test. 15

Supp. Fig. 9: Joint test power and calibration for other trait pairs. 16

Supp. Fig. 10: Joint estimates under complementary selection. 17

Supp. Fig. 11: Joint test, including/excluding the causal trait. 18

Supp. Fig. 12: Correlated response test, including/excluding the causal trait. 19

Supp. Fig. 13: K-way tests for selection and correlated response. 20

References 21

### Appendix

#### Inference

##### Importance sampling estimation of the likelihood function of selection

Our likelihood model builds heavily on our previous work, which developed importance sampling approaches to estimating the likelihood function of the selection coefficient acting on a SNP, $L^{SNP}(s)$.[1] Here, we briefly explain the importance sampling method used to estimate $L(\omega)$, the likelihood of the multivariate selection gradient:

$$L(\omega) = \prod_{i=1}^{M} \quad L_i^{SNP}\left(\beta_{(i)}^{\intercal}\omega\right), \tag{A1}$$

where $\beta_{(i)}$ is the vector of trait effects for SNP $i$. In the following, we omit the subscript $i$ for brevity. We can model the relationship between SNP $s$ and the haplotype data $D$ from a window around the SNP via the latent ancestral recombination graph (ARG) $G$,

$$L^{SNP}(s) = E_p[P(D \mid G, s)] = E_q\left[P(D \mid G, s)\frac{p(G \mid s)}{q(G)}\right] \tag{A2}$$

for any appropriate choice of $q$ such that $p(s) > 0 \Rightarrow q(G) > 0$, which generally will hold in our case. Thus, we can approximate the SNP likelihood function as

$$\hat{L}^{SNP}(s) \coloneqq \frac{1}{m}\sum_{l=1}^{m} \quad P\left(D|G^{(l)}, s\right)\frac{p(G^{(l)} \mid s)}{q(G^{(i)})} \rightarrow L_k^{SNP}(s) \tag{A3}$$

Where the convergence is almost surely as $m \rightarrow \infty$.

We are interested in the particular choice of $q(G) = p(G \mid D, s = 0)$, the posterior under selective neutrality, because programs such as *ARGweaver*[2] and *Relate*[3] can be used to approximately sample the posterior ARG, or aspects of it (e.g. a local tree). We showed previously that the approximation

$$\widehat{LR}^{SNP}(s) = \frac{1}{m}\sum_{l=1}^{m} \quad \frac{p\left(G_i^{(l)} \mid s\right)}{p\left(G_i^{(l)} \mid s=0\right)} \tag{A4}$$

is a tractable and accurate estimate of the likelihood ratio of $s$, where $G_i$ denotes the local tree at SNP $i$, extracted from the ARG $G$. Here, we introduce and use a slightly different estimator,

$$\widehat{LR}^{SNP}(s) = \frac{\sum_{l=1}^{m} \quad \frac{p\left(G_i^{(l)} \mid s\right)}{\pi\left(G_i^{(l)}\right)}}{\sum_{l=1}^{m} \quad \frac{p\left(G_i^{(l)} \mid s=0\right)}{\pi\left(G_i^{(l)}\right)}} \tag{A5}$$

where $\pi(\cdot)$ is a neutral prior on coalescence trees. While $p(\cdot)$ is calculated using the structured coalescent, with lineages subtending the same allele with frequency $X(t)$ coalescing at rate $\lambda(t) = N(0)/[N(t)X(t)]$, the prior $\pi(\cdot)$ is calculated using the unstructured coalescent with rate $\lambda(t) = N(0)/N(t)$. Note that we do not explicitly model population structure (e.g. gene flow).

We also note that we have made several additional modifications to the importance sampling approximation of the likelihood ratio: first, we assume that the allele frequency trajectory is a deterministic, logistic function of time, when previously we modeled stochasticity in the allele frequency trajectory (see the next section for more details). Because we focus on applying our method to detecting adaptation in the recent past, this approximation is appropriate when drift has had little opportunity to distort allele frequencies.

Second, we make a functional approximation to $\log\widehat{LR}^{SNP}(s)$. We do a grid search for the optimal value of $s^*$, and then we fit a quadratic function to points $\{(s, log\ \widehat{LR}^{SNP}(s)) : |s - s^*| < \delta\}$. Optimizing $\log\widehat{LR}(\omega)$then becomes a simple process of solving a linear system of equations:

$$\log \widehat{LR}(\omega) = \sum_i \quad (a_i(\beta_{(i)}^{\intercal}\omega)^2 + b_i(\beta_{(i)}^{\intercal}\omega) + c_i) \qquad \text{(A6)}$$

Where $(a_i, b_i, c_i)$are the fitting coefficients of the quadratic approximation for SNP $i$, in descending order of degree. Thus

$$\widehat{\omega} = [2\sum_i \quad a_i\beta_{(i)}\beta_{(i)}^{\intercal}]^{-1}(\sum_i \quad b_i\beta_{(i)}). \qquad \text{(A7)}$$

This approximation has two benefits: (1) solving for the selection gradient estimate is extremely simple and fast, and (2) it makes it feasible to calculate standard errors using resampling approaches.

#### Accounting for multiple SNPs in LD

In our analyses we assume independence of local LD blocks (see e.g. Berisa & Pickrell, 2016). Generally we choose to ascertain a single SNP for each LD block and include its SNP likelihood in the product (Eq. A1). However, in joint analyses it may be necessary to ascertain multiple SNPs per LD block, each corresponding to a GWAS hit for a different trait.

Let $B(i)$ denote the set of ascertained SNPs in the same LD block as $i$. If only 1 SNP from each LD block is included, then $B(i) \quad = 1$for each ascertained SNP $i$. If multiple SNPs from the same LD block are included, we exponentiate each of these SNPs' likelihoods by a factor $1/|B_i|$:

$$L(\omega) = \prod_{i=1}^{M} L_i^{SNP}(\beta_{(i)}^{\intercal}\omega)^{1/|B_i|} \qquad \text{(A8)}$$

This can be considered a conservative method for dealing with SNPs in LD. For example, let $A$be our set of ascertained SNPs. If two nearby SNPs $i_1, i_2$ are in perfect LD ($r^2 = 1$), then we expect $L_{i_1}^{SNP}(s)$=$L_{i_1}^{SNP}(s)$ and

$\beta_{(i_2)}$=$\beta_{(i_2)}$. Suppose all other SNPs in $A$are independent (i.e. ascertained from distinct LD blocks). Then the exponentiation factor recovers the original likelihood

$$L(\omega) = \prod_{i=1}^{M} L_i^{SNP}\left(\beta_{(i)}^{\intercal}\omega\right)^{1/|B_i|}$$

$$= \sqrt{L_{i_1}^{SNP}(s) \cdot L_{i_2}^{SNP}(s)} \cdot \prod_{i \in S:\, i \neq i_1, i_2} L_i^{SNP}\left(\beta_{(i)}^{\intercal}\omega\right)$$

$$= \prod_{i \in S:\, i \neq i_2} L_i^{SNP}\left(\beta_{(i)}^{\intercal}\omega\right) \quad = \prod_{i \in S:\, i \neq i_1} L_i^{SNP}\left(\beta_{(i)}^{\intercal}\omega\right) \qquad \text{(A9)}$$

In the other limiting case $r^2 = 0$, this correction factor is conservative, as it discounts the contribution of $i_1, i_2$ to the log likelihood by a factor of ½.

##### Selection gradient and correlated selection standard errors

We use a block-bootstrap approach to calculating the standard errors of $\hat{\omega}$. Specifically, we identify LD blocks and bootstrap loci ascertained in distinct blocks. Given the standard errors, we assess significance using a Wald test on the $Z$ statistic $\hat{\omega}/\widehat{se_{\omega}}$.

We also compute a statistic we call $R$ to assess whether a trait $j$ has evolved under correlated response to selection on some disjoint set of traits $T$. To do this, we can estimate selection gradients for two sets of traits, $T$and $T \cup \{j\}$, and calculate $R \;\; = \omega^{(T \cup \{j\})} - \omega^{(\{j\})}$ where $\omega^{(U)}$is the selection gradient of the trait estimated with respect to a set of traits $U$, calculate $\widehat{se_R}$ through block-bootstrap, and assess significant using a Wald test on $R \;\; / \widehat{se_R}$ .

### Coalescent likelihood models

##### *Relate* prior

The prior $\pi(T)$is the standard coalescent with changing effective population size. First, let $U$be the vector of $n - 1$ coalescent times of $T$, ordered most to least recently. Due to exchangeability of lineages, the density only depends on $T$ via these coalescent times $U$. Specifically,

$$\pi(T) = \prod_{i=1}^{n-1} \quad p(U_i = u_i \mid U_{i-1} = u_{i-1}) \qquad \text{(A10)}$$

$$p(U_i = u_i \mid U_{i-1} = u_{i-1}) = \frac{n-i+1}{2} \cdot N(0)/N(u_i) \cdot exp\left(-\frac{n-i+1}{2}(\Lambda(u_i) - \Lambda(u_{i-1}))\right) \qquad \text{(A11)}$$

$$\Lambda(u) = \int_0^u \quad N(0)/N(t) \;\cdot dt \qquad \text{(A12)}$$

We assume that $N(t)$ is piecewise constant and can be expressed using $\tau = (\tau_0, \tau_1, \dots)$ and $N = (N_0, N_1, \dots)$ such as the required models for *ARGweaver* and *Relate;* hence, finding $\Lambda(u)$ is a simple sum over integrals defined over constant functions:

$$\Lambda_i = \sum_{k=1}^{b(u_i)} N_0 \tau_k / N_k + N_0 (u_i - \tau_{b(u_i)}) / N_{b(u_i)} \tag{A13}$$

where $b(u) := max \{k \in (0,1,2 \dots) : u > \tau_k\}$.

#### Coalescent selection likelihood under deterministic model

Unlike in our previous work,[1] in which we treated the allele frequency as a latent random variable, here we use a deterministic approximation of the allele frequency trajectory. Under the standard 'hard sweep' model, an appropriate approximation would be $X(t \mid s) = (1 + (1 - x_0)/x_0 \cdot e^{st})^{-1}$. Technically, if we want to express the trajectory conditional on the present-day derived allele frequency (DAF) $x_0$, it would be more appropriate to use a closer approximation of the backwards Wright-Fisher diffusion with selection (see e.g. [4]). However, since we are mostly interested in modeling the recent past for common alleles ascertained in a GWAS (usually DAF > 1%), this approximation is appropriate, especially in populations of large recent $N_e$ such as humans, where drift is negligible on short timescales.

We assume a pulse of selection over some time interval $(a, b)$, outside of which the allele is effectively neutral (and, we assume, at constant frequency):

$$X(t, s, x_0) = x_0 , \quad t < a \tag{A14}$$

$$= (1 + (1 - x_0)/x_0 \cdot e^{s(t-a)})^{-1} , \quad a \le t < b \tag{A15}$$

$$= (1 + (1 - x_0)/x_0 \cdot e^{sb})^{-1} , \quad t > b \tag{A16}$$

To calculate $p(T \mid s)$, we split the tree into two subtrees (imagine 'deleting' the branch on which the mutant allele arose). Note that we implicitly assume the site is biallelic, such as under the infinite sites assumption. Let us label these alleles $A1$ and $A2$; these labels must be consistent with the polarization of the GWAS summary statistics; we assume that those are polarized w.r.t the $A1$ allele. Within each of these subtrees, we find the coalescent times $U^{A1}$ and $U^{A2}$. Then

$$p(T \mid s) = \prod_{i=1}^{n_1 - 1} p \left(U_{n-i} = u_i^{A1} \mid U_{n-i+1} = u_{i-1}^{A1}, s, x_0\right) \times$$

$$\times \prod_{i=1}^{n_2-1} p(U_{n-i} = u_i^{A2} \mid U_{n-i+1} = u_{i-1}^{A2}, -s, 1 - x_0) \tag{A17}$$

$$p(U_{k-1} = t \mid U_k = t', s, f) = \frac{k}{2} \cdot \frac{N(0)}{N(t)X(t)} \cdot exp \left(-\frac{k}{2}(\Lambda(t, s, f) - \Lambda(t', s, f))\right)$$

(A18)

$$\Lambda(t,s,f) = \int_0^t \quad N(0)/[N(\tau)X(\tau,s,f)] \cdot d\tau \tag{A19}$$

where $U^{A1}$ and $U^{A2}$ are measured in units of $2N(0)$ generations.

### Supplementary Figures

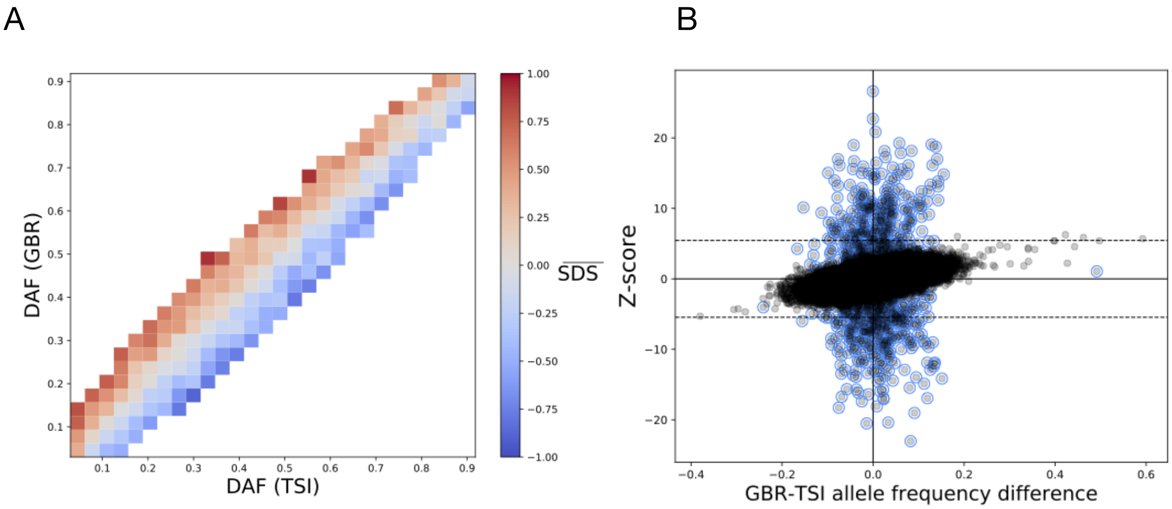


**Supp. Fig. 1: Distribution of frequencies and SDS in 1000 Genomes SNP set.**
(A) mean SDS score with respect to joint derived allele frequency (DAF) in TSI *vs* GBR. We selected a set of 40,320 autosomal SNPs with MAF >0.5% in the UK Biobank and $MAC_{GBR+TSI} \geq 4$, HWE $P > 10^{-6}$in the 1000 Genomes Phase 3 data, which we used in our simulations of uncorrected stratification. We found that this SNP set recapitulates the pattern demonstrated in Sohail, *et al.* (2019); namely, that SNPs with higher frequency in GBR tend to have higher SDS, and vise versa for TSI.[5] DAF was calculated from 1000 Genomes phase 3 data for all autosomes and SDS was obtained from previous analysis of the UK10K cohort.[6] To limit noise, we show DAF bins with ≥30 SNPs. (B) An example simulation of uncorrected stratification in Z-scores of the aforementioned SNP set. Here we set $h^2 = 50\%, M = 10^3, N = 10^5, N_{TSI}/N_{GBR} = 5\%, \sigma_S = 0.1$. Causal SNPs are circled in blue. Dashed lines indicate genome-wide significance thresholds ($P < 5 \times 10^{-8}$).

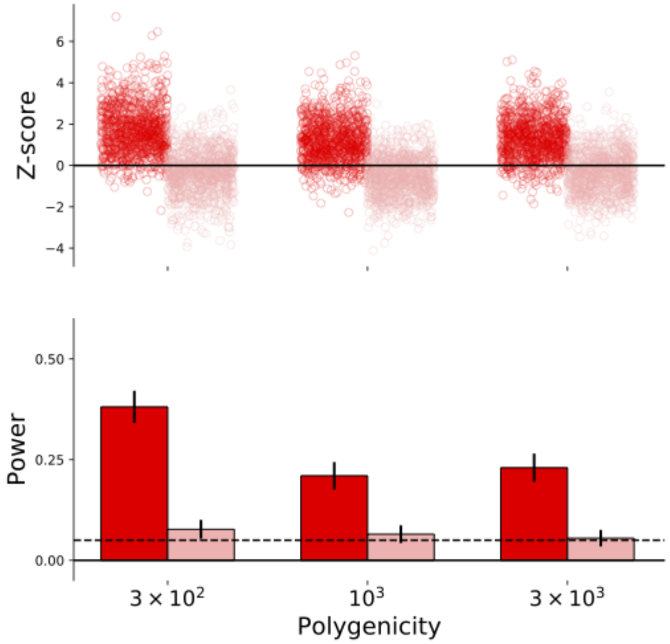


**Supp. Fig. 2: Calibration and power under GBR demography.**
We simulated polygenic adaptation under a model of changing population size based on GBR 1000 Genomes individuals[3] with $\omega = 0.05$, and a pulse of recent selection over the last 35 generations. Red denotes simulations with selection, pink denotes neutral simulations. Dashed lines indicate nominal FPR (5%) and black lines denote 95% Bonferroni-corrected CIs. We simulate a trait with $h^2 = 50\%$and $N = 10^5$. To estimate trees we used a sample size of $n = 400$ haplotypes of length 1Mb and assume mutation and recombination rates of $\mu = r = 10^{-8}$/bp/gen. We simulate 1000 replicates in each case.

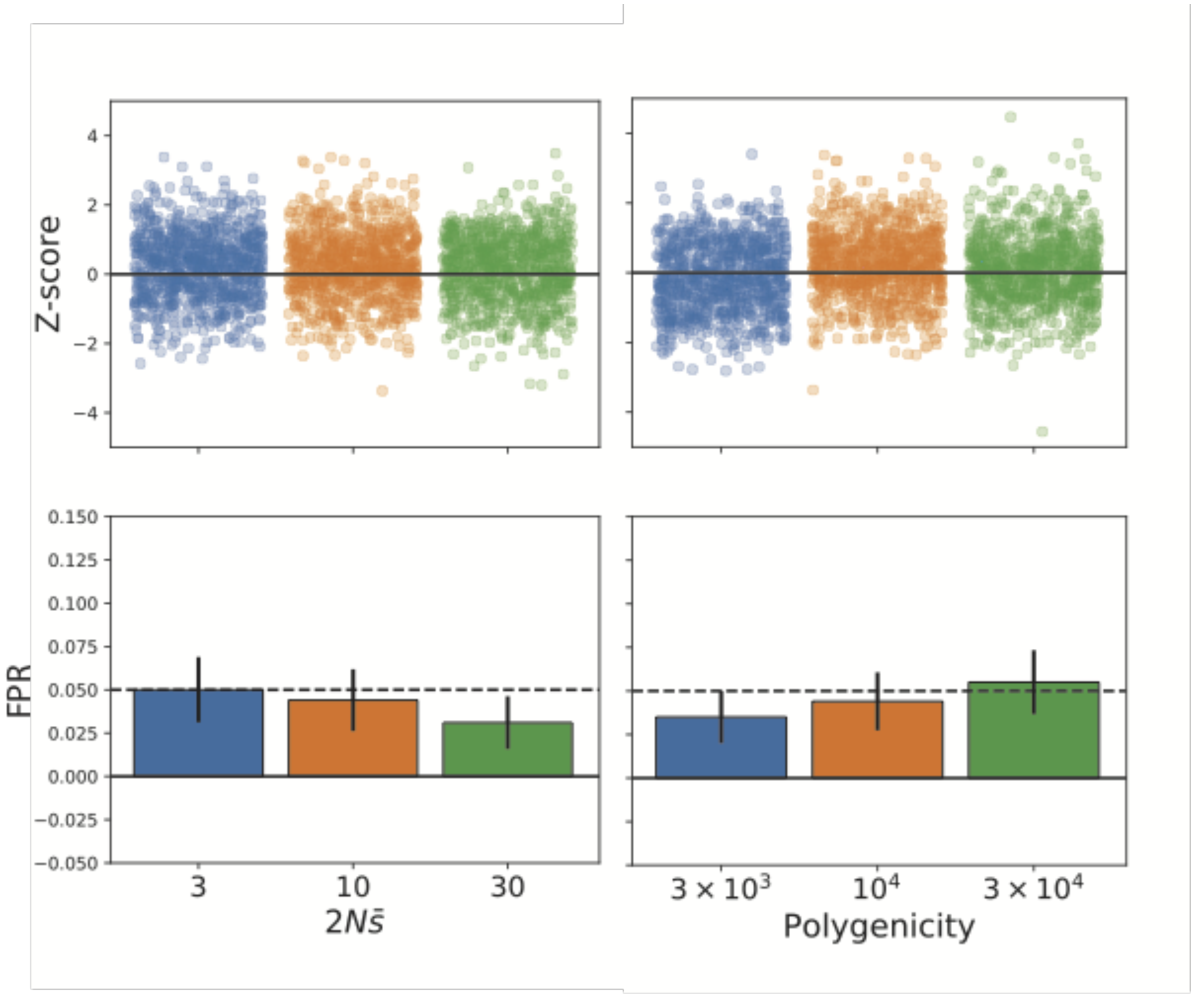


**Supp. Fig. 3: Robustness to purifying selection.**
We simulated traits under purifying selection using the model of Schoech, *et al.* (2019).[7] That is, we model the joint distribution of allele frequency and selection coefficient of a causal SNP. We assume that SNPs are always deleterious and the magnitude of the selection coefficient has an exponential distribution, with mean $\bar{s}$. Given $s$ for a particular SNP, the allele frequency is drawn randomly from its stationary distribution. We assume that $|\beta| = c \cdot s^{1/2}$ -- the constant of proportionality is chosen post-hoc to normalize SNP heritabillity to 50% -- so our results also approximate for the dynamics of Gaussian stabilizing selection, modulo underdominance and epistatic effects. Dashed lines indicate nominal FPR (5%) and black lines denote 95% Bonferroni-corrected CIs. We simulate GWAS with $N = 10^5$. To estimate trees we used a sample size of $n = 400$haplotypes of length 1Mb and assume mutation and recombination rates of $\mu = r = 10^{-8}$/bp/gen. We simulate 1000 replicates in each case.

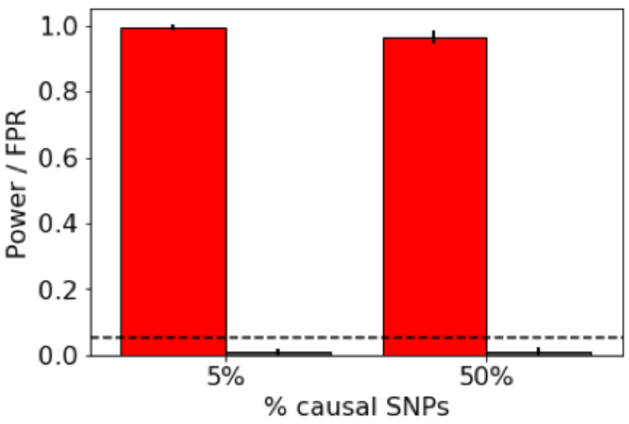


**Supp. Fig. 4: Calibration and power under allelic heterogeneity.**
We simulated polygenic adaptation of a trait with multiple linked causal SNPs in LD (i.e., allelic heterogeneity) in SLiM[8]. We assume $h^2 = 50\%$and a mutational target of 100×100kb regions with $\mu = r = 10^{-8}$. We assume a constant population size of $N_e = 10^3$ and burn in simulations for 9990 generations, and then for the next 10 generations, we simulate a directional selection gradient of $\omega = 0.2$ (we chose this value because it resulted in ~1SD increase in the mean phenotype). We simulated two levels of allelic heterogeneity; (left) 5% and (right) 50% of mutations in the target are causal. Red bar indicate power to detect simulations with selection; pink bars (which are too short to see this color) indicate FPR under the null, $\omega = 0$. SNPs were ascertained by taking one at each independent regions with the maximum value of $2pq\beta^2$within the region. We tested for selection using the true local trees at these sites. Dashed lines indicate nominal FPR (5%) and black lines denote 95% Bonferroni-corrected CIs. We simulated 1000 replicates in each case.

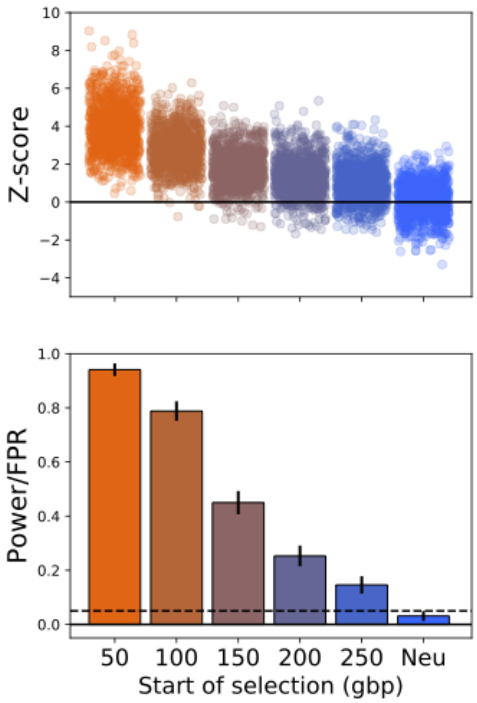


**Supp. Fig. 5: Time specificity of test for recent selection.**

We simulated under models of different timings of selection, assuming a 50-generation pulse with start time ranging from 50 to 250 generations ago, as well as neutral simulations, and run PALM under a nominal model of selection in the last 50 generations. Dashed lines indicate nominal FPR (5%) and black lines denote 95% Bonferroni-corrected CIs. We simulate GWAS with $N = 10^5$. To estimate trees we used a sample size of $n = 400$haplotypes of length 1Mb and assume mutation and recombination rates of $\mu = r = 10^{-8}$/bp/gen. We simulate 1000 replicates in each case.

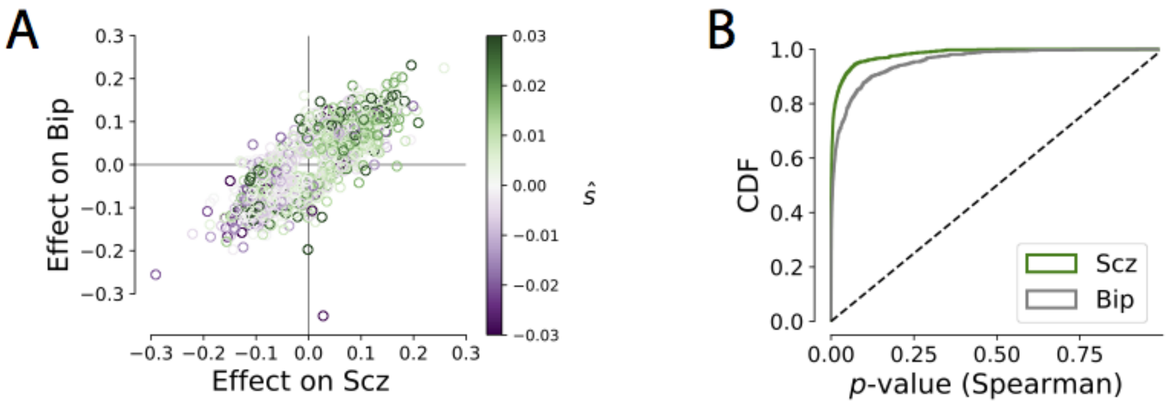


**Supp. Fig. 6: Pleiotropy causes bias in tests for polygenic adaptation.**
We simulated a bivariate trait (modeled after $h^2$and $r_g$of schizophrenia [Scz], and bipolar disorder [Bip]). We simulated a pulse of selection acting to increase Scz prevalence over the last 50 generations. (A) Estimates of directional selection on SNPs ($\hat{s}$) are positively correlated with both SNP effects for Scz and Bip. We estimated selection using our importance sampling method and included SNPs ascertained in our simulated GWAS. (B) Testing for selection on a neutral correlated trait (here, Bip) yields massive inflation of the false positive rate. We evaluated p-values by computing significance of the Spearman correlation of $\hat{\beta}$ and $\hat{s}$.

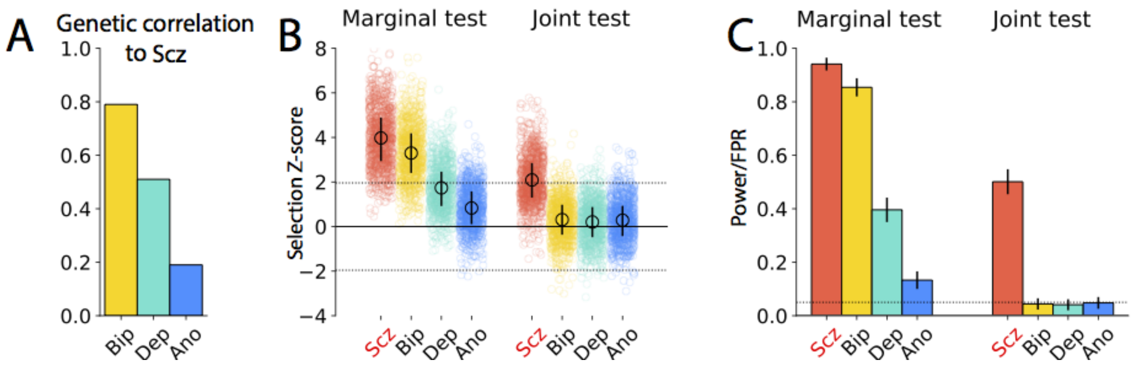


**Supp. Fig. 7: Marginal vs. joint test comparison, lower pleiotropy.**
Here we recreate Figure 3 (main text) but setting the level of pleiotropy used to simulate the polygenic trait architectures to be lower ($\varrho$ = *60%*). See Figure 3 for all simulation details.

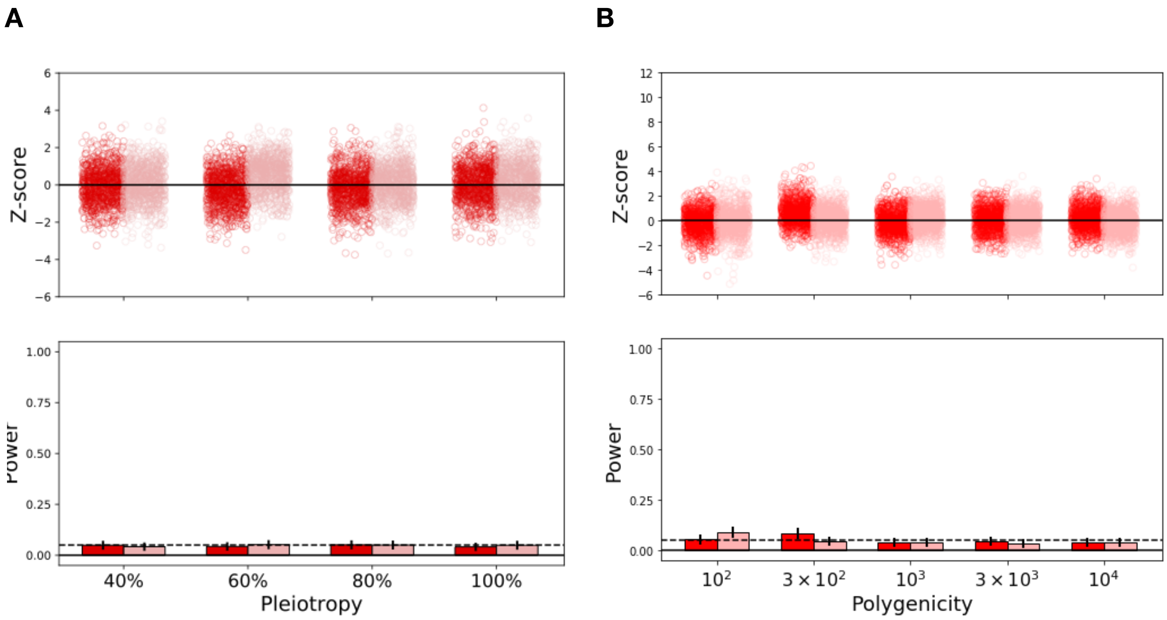


**Supp. Fig. 8: Calibration of joint test.**

Simulations under neutrality, testing Trait I and Trait III jointly. Here we show calibration of J-PALM when neither of the two traits is under selection (i.e., both are neutral). We consider the effects of degree of pleiotropy (A) and polygenicity (B). Other simulation details are identical to Figure 3.

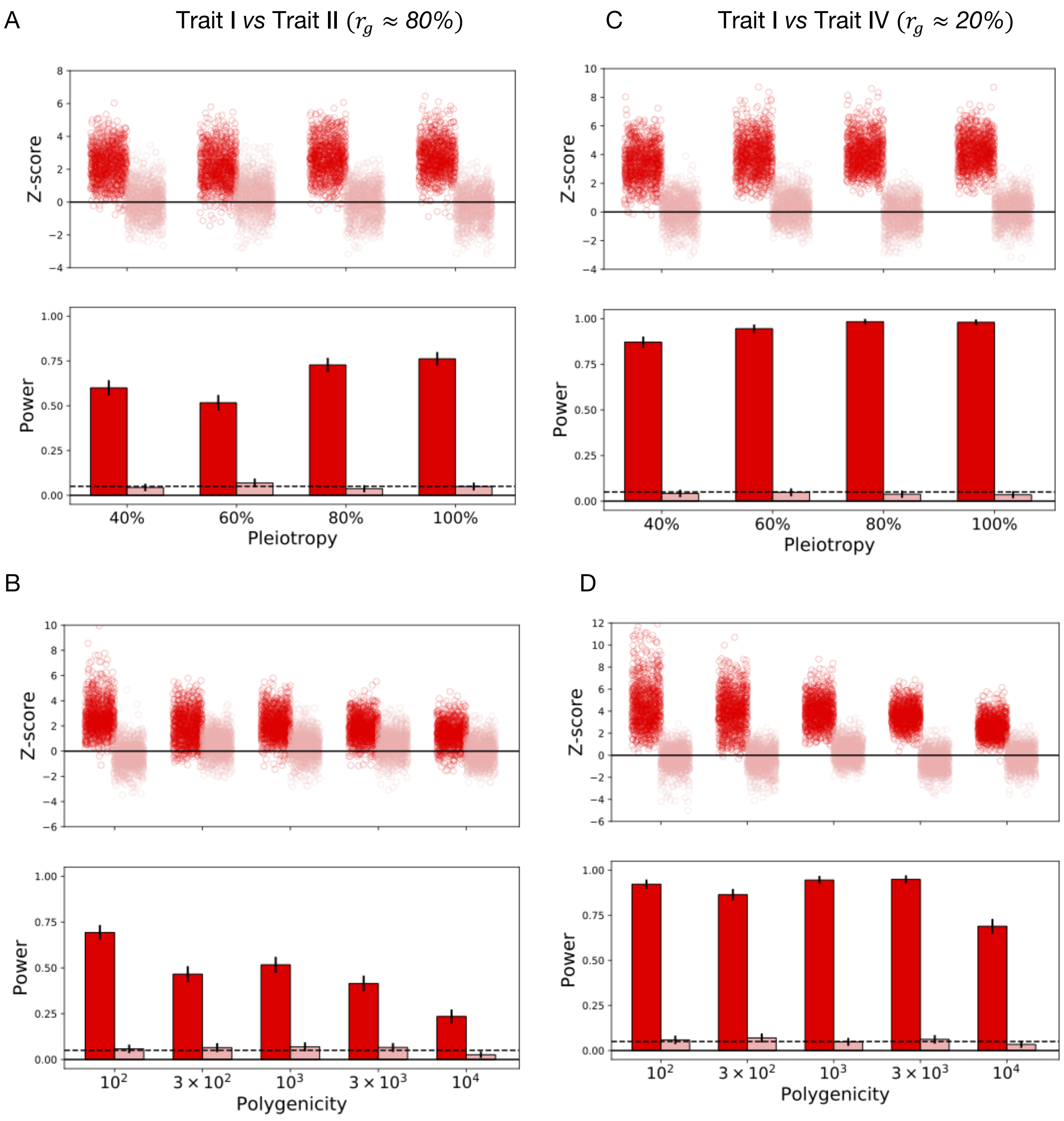


**Supp. Fig. 9: Joint test power and calibration for other trait pairs.**
Simulations under neutrality, using J-PALM to test Trait I *vs* Trait II (A,B) and Trait I *vs* Trait IV (C,D) jointly. We varied degree of pleiotropy $\varrho$ (A,C) and polygenicity $M$ (B,D). In all simulations we simulated selection to increase Trait I, with all other traits neutral. Other simulation details are identical to Figure 3.

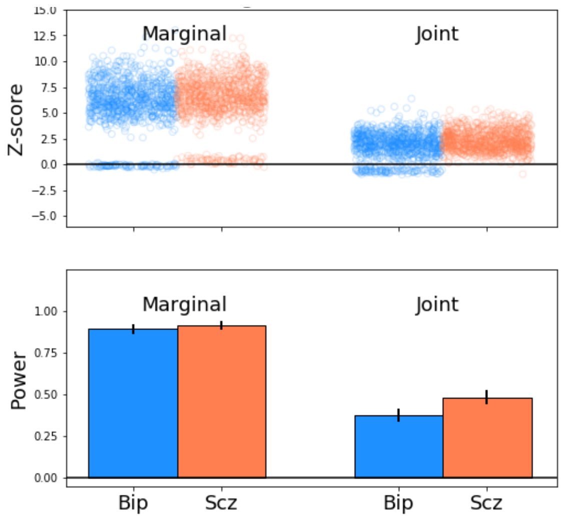


**Supp. Fig. 10: Joint estimates under complementary selection.**
Positive selection ($\omega$ = *0.05*) simulated on both of two traits, Trait I & II, modeled after SNP heritability and genetic correlation of bipolar (Bip) and schizophrenia (Scz). Selection was estimated marginally (left) and jointly (right). Simulations follow the approach demonstrated in Fig. 3.

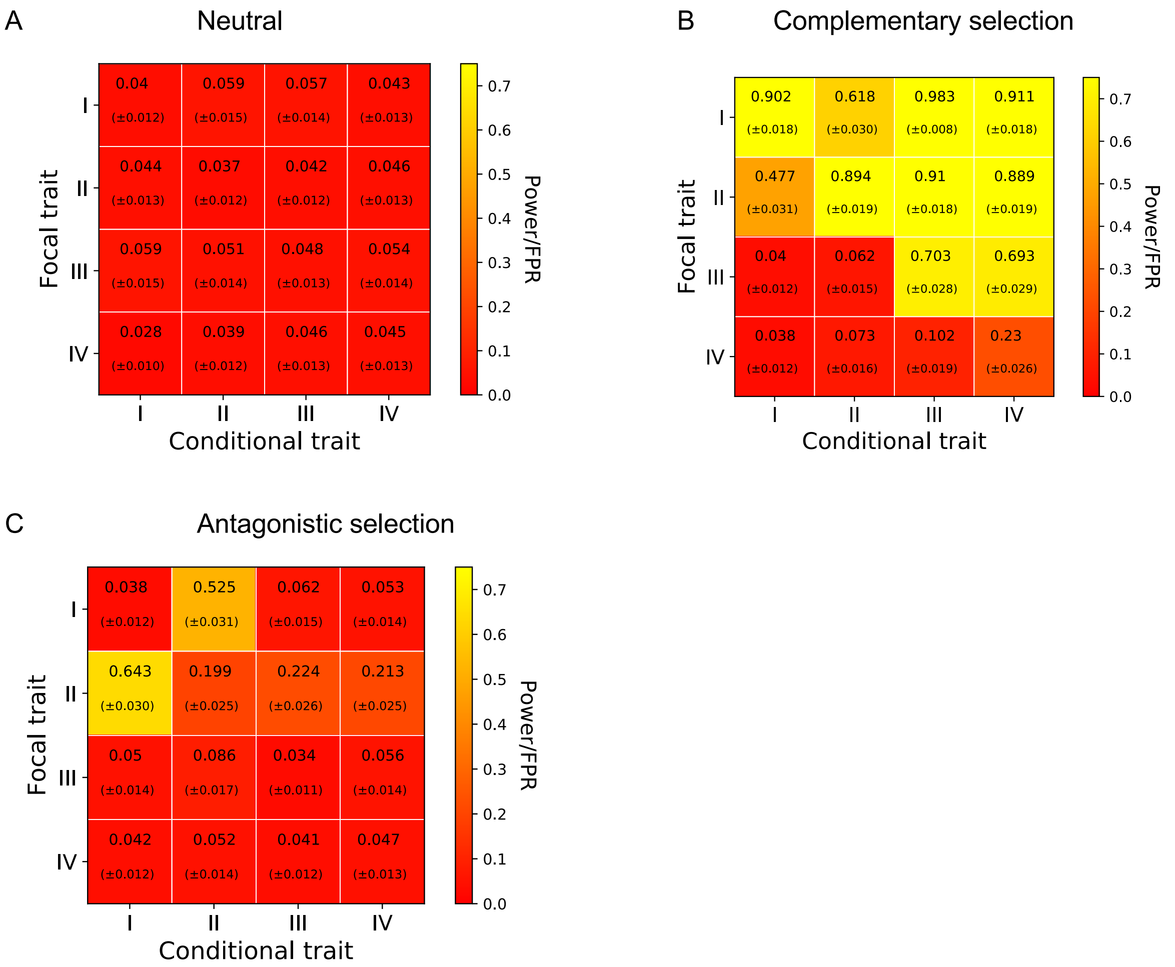


**Supp. Fig. 11: Joint test, including/excluding the causal trait.**
We conducted joint tests over all pairs of Traits I/II/III/IV under 3 scenarios: (A) neutral (all traits neutral), (B) Traits I & II under positive selection, (C) Trait I under positive selection, Trait II under negative selection. Text and color show positive rate. Parentheses indicate standard errors.

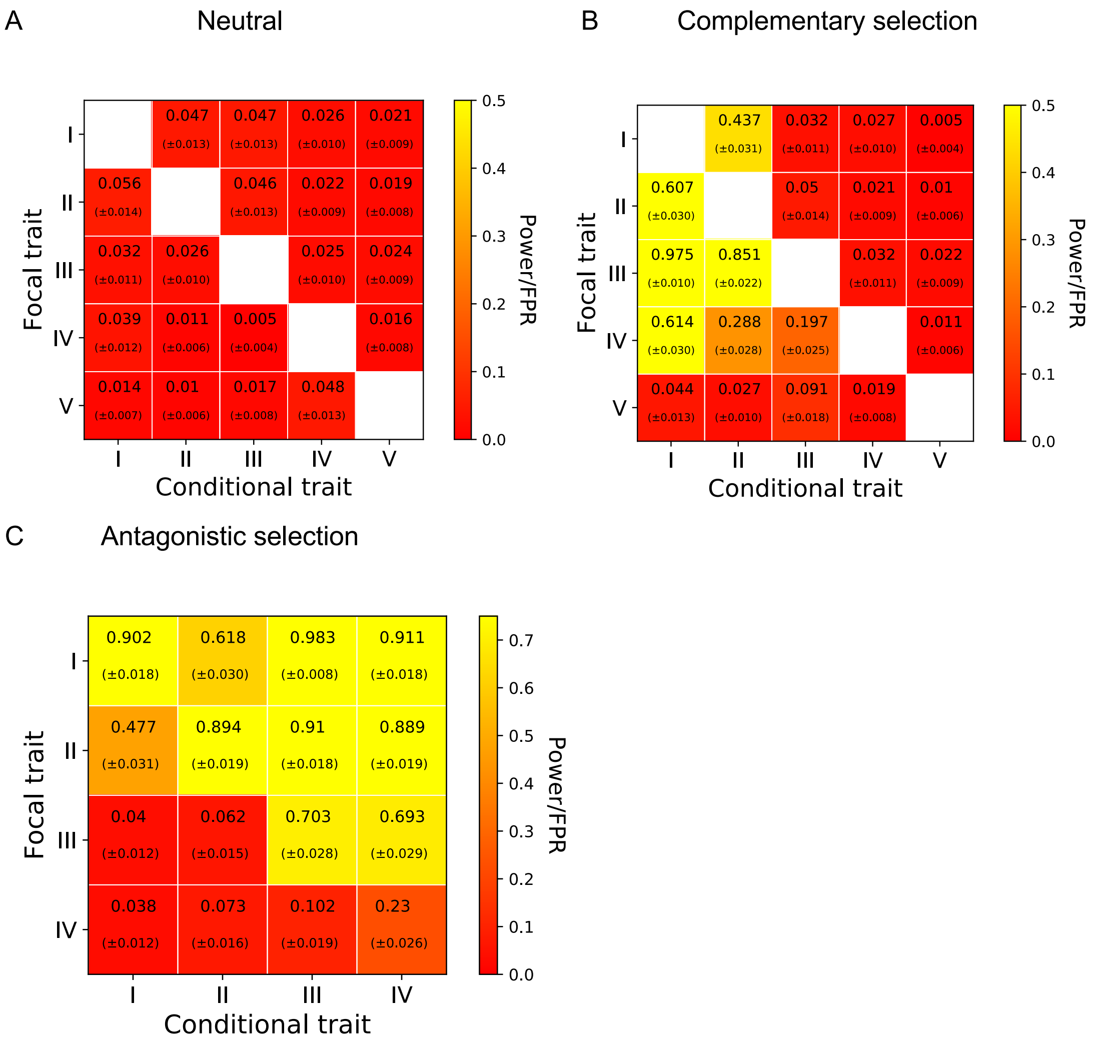


**Supp. Fig. 12: Correlated response test, including/excluding the causal trait.**
We conducted tests for correlated response over all pairs of Traits I/II/III/IV under 3 simulation scenarios: (A) neutral (all traits neutral), (B) Traits I & II under positive selection, (C) Trait I under positive selection, Trait II under negative selection. Text and color show positive rate. Parentheses indicate standard errors.

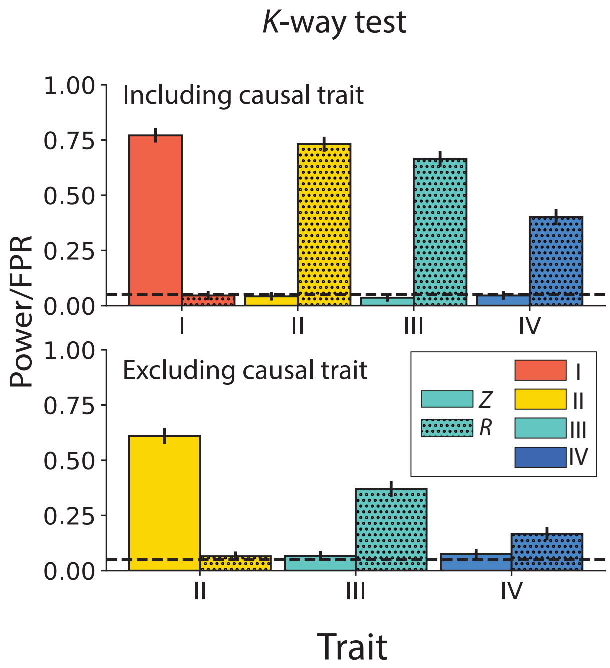


**Supp. Fig. 13: K-way tests for selection and correlated response.**

Top: including all traits. Bottom: excluding trait I. Solid bars: test positive rate for selection test. Dotted bars: test positive rate for correlated response test. Trait I is under positive selection, with all other traits neutral.
